# Optimized cryo-FIB milling strategy to generate thin, minimally damaged biological lamellae

**DOI:** 10.64898/2026.07.21.739890

**Authors:** Laina N. Hall, Joshua Paul, Philip Ngo, Bronwyn A. Lucas

## Abstract

Focused ion beam (FIB)-milling has been adapted to thin frozen cells for visualization of macromolecular structures *in situ* with cryogenic electron microscopy. However, only a few large and abundant complexes have been annotated to date. FIB-milling introduces damage which limits the recoverable information from cellular sections. Here, we present Nilas, a low-energy milling strategy optimized to minimize damage and produce thin lamellae. Nilas-milled lamellae show minimal FIB-milling damage, contain areas at or below 50 nm and produce higher resolution *in situ* 3D reconstructions. Nilas improves the recovery of ribosomal subunits and reduces the predicted minimal detectable molecular mass with two-dimensional template matching (2DTM) to ~220 kDa. Consistently, we recover additional non-ribosomal complexes including RNA polymerase III with 2DTM in Nilas-milled lamellae. Nilas is compatible with common milling hardware, making it accessible to diverse users. By extending the size limit for *in situ* structural biology we bring visual proteomics closer to reality.

## Introduction

Cryogenic electron tomography (cryo-ET) and microscopy (cryo-EM) are powerful tools to visualize macromolecules in their native cellular environment (*in situ*) ^1^. However, most cells are too thick to image directly with cryo-EM. Cryogenic focused ion beam (FIB)-milling is the main method to thin frozen cells to imageable sections (lamellae) with improved preservation of cellular ultrastructure relative to current cryogenic ultramicrotomy protocols ^2–6^.

Cellular lamellae generated by cryo-FIB milling are typically 2–6 times thicker than single particle samples resulting in a ~15–40% lower signal-to-noise ratio (SNR). The lower SNR limits the achievable resolution of *in situ* structures and prevents the detection of particles. Relative to single particle analysis, the reduced SNR in cellular samples corresponds to a hypothetical increase in minimal alignable molecular mass of ~30–200% ^7,8^. Decreasing the thickness of cellular lamellae has the potential to maximize the SNR, improve the resolution of *in situ* structures and increase the detectable proportion of the proteome.

However, the potential benefits of producing thinner lamellae are limited by FIB-milling damage. In biological lamellae, FIB-milling introduces damage which decays exponentially from the surface and can be detected at depths up to 40–60 nm depending on the method used to measure the damage ^9–13^. The optimal lamella thickness is therefore a balance between electron scattering, which reduces SNR in thicker samples, and FIB-milling damage which becomes dominant in thinner samples. With a gallium liquid metal ion source (LMIS) FIB and standard milling strategies, this balance is achieved at ~90-100 nm ^9^. In thinner lamellae, damage causes signal to degrade ^9^. Consistently, the thinnest lamellae do not produce the highest resolution reconstructions ^14,15^. In lamellae of typical thickness, 150–250 nm, FIB-milling damage reduces the recoverable signal by 10–25% ^9^. Therefore, minimizing FIB-milling damage would improve the recovery of signal from thicker lamellae and capitalize on the improved SNR that comes from thinning below 100 nm.

Optimizing FIB-milling for visual proteomics requires a rapid and sensitive assay for the integrity of particles within the lamella. We and others have demonstrated that two-dimensional template matching (2DTM) can be used to localize specific macromolecules in untilted micrographs by cross-correlation with a high-resolution reference template ^7,16,17^. The use of untilted micrographs in place of a tilt series dramatically reduces the time required for data collection and maximizes the recovery of high-resolution features ^18^. 2DTM produces a pixel-wise z-score which is used to identify significant detections by comparison to a threshold set to minimize false positives. The 2DTM z-score is sensitive to small deviations between the reference template and target, including particle damage from FIB-milling. By varying the contrast transfer function applied to the template, 2DTM can be used to estimate the particle *z*-position to within 2 nm, providing a depth dependent metric of particle quality ^7,9,16,17,19^. We have previously established a sensitive, single molecule assay that uses changes in the 2DTM z-score to quantify FIB damage ^9^. In this study we leverage this 2DTM-based assay to identify optimized milling conditions that maximize signal recovery from cellular lamellae.

FIB-milling damage results from ion implantation, secondary collisions and radiation damage from secondary electron generation ^20^. Lowering the accelerating voltage reduces the depth of ion implantation and the range of secondary collisions and electrons ^21^. Reducing the accelerating voltage from 30 kV to 8 kV during the final step of lamella preparation reduced damage in lamellae milled from purified virus samples and plunge frozen cells from 50–60 nm to 30 nm ^12,22^. Voltages lower than 8 kV would further reduce ion-beam damage but have not been applied to cellular lamellae because the unfocused beam is not precise enough for standard polishing protocols. Alternate low voltage strategies have been developed for the generation of thin, undamaged lamellae for atomic-resolution scanning transmission electron microscopy (STEM) using both gallium and plasma FIBs^23,24^, but have not yet been adapted for generating biological lamellae.

Here, we present Nilas, a FIB-milling strategy to generate thin and minimally damaged cellular lamellae. Nilas accounts for the less focused beam at low voltages and maximizes overall milling speed by minimizing the volume polished at low voltages. Specifically, we use three voltages, reducing from 30 kV to 8 kV to a final 2 kV polish with a 10 degree over-and under-tilt. Nilas reduces FIB-milling damage to close to the the limit of detection and enables the production of lamellae with regions of 50 nm and below. Images collected from Nilas-milled lamellae have a higher SNR, resulting in higher resolution 3D reconstructions and the reduction of the estimated minimal detectable molecular mass with 2DTM to ~220 kDa. Nilas improves the detection of diverse macromolecular complexes with 2DTM including small and large ribosomal subunits, the 20S proteasome, RNA polymerase III and glycolytic enzymes in Nilas-milled lamellae.

## Results

### Determining optimal milling conditions to generate thin, minimally damaged lamellae

We have previously identified FIB-milling damage as a major factor limiting the retrievable signal from biological lamellae and preventing potential improvements from thin lamellae. To develop a strategy to reliably generate thin, undamaged lamellae, we first explored milling with different ions. We compared lamellae milled with a gallium FIB generated from a liquid metal ion source (LMIS) and argon and xenon FIBs generated from plasma, using standard milling protocols (Supplementary Fig. S1a). We used 2DTM to identify large ribosomal subunits (LSUs) in images containing cytoplasm (Supplementary Fig. S1b) (PDB: 6Q8Y) ^25^. The change in 2DTM z-scores as a function of distance from the lamella surface was used as a single molecule damage metric ^9^. Consistent with our prior work, we find that the 2DTM z-scores decrease towards the lamella surface in images of 30 kV-milled lamellae (Supplementary Fig. S1c). Lamellae milled with xenon were significantly (*p<*0.05, t-test) less damaged relative to lamellae milled with gallium or argon when operated at 30 kV (Supplementary Fig. S1d). More large ribosomal subunits were detected toward the lamella surface in xenon-milled lamellae (Supplementary Fig. S1e). Despite this improvement, we observed considerable damage detectable to a depth of 50 nm with all ions at 30 kV (Supplementary Fig. S1).

To further reduce milling damage, we explored milling at lower accelerating voltages. We chose to polish with 2 kV because lower voltages (500 V, 1 kV) show minimal differences in predicted ion implantation relative to 2 kV (Supplementary Fig. S2a) but greatly increase the predicted probe size (Supplementary Fig. S2b,c). As an additional consideration, 2 kV is the lowest accelerating voltage available in the most common gallium cryo-FIB SEM hardware, namely the Aquilos (Thermo Fisher) and Crossbeam (Zeiss) increasing the accessibility to more users. We chose to use a gallium LMIS FIB for three main reasons. Firstly, at 2 kV, gallium is better focused than xenon at the low currents used for polishing (Supplementary Fig. S2c). Secondly, the difference in the predicted median implantation range between gallium, argon and xenon becomes negligible at 2 kV (Supplementary Fig. S2a,d-f) ^26^. Thirdly, sputter rates of xenon and gallium are similar at 2 kV ^27^. Therefore, although xenon can be operated at higher currents and produced less damaged lamellae at 30 kV, we do not expect to see a difference in milling speed or damage when polishing at 2 kV.

The diffuse beam at 2 kV also presents challenges for imaging and milling lamellae. At 2 kV, there is sufficient contrast to reliably locate lamellae but not to accurately place a milling area for polishing (Supplementary Fig. S2g-i).

Standard biological cryo-FIB milling protocols require a focused beam to precisely polish parallel to the lamella surface and are therefore incompatible with a broad, low-voltage beam. A voltage step-down with high tilts (up to ±7°) during low-voltage (*<*2 kV) polishing was previously developed to produce thin, undamaged silicon samples for atomic-resolution STEM^23^. We adapted this approach to mill cryogenic biological lamellae. To reduce lamella breakage and curtaining during later, low-voltage steps we increased the organo-platinum layer deposited with a gas injection system (GIS) by two-fold and performed 30 kV rough milling using an over-tilt and under-tilt of ±1.5°(Fig. 1a). To account for the broader beam at 8 kV we increase the distance between the cleaning cross section (CCS) box and the lamella and used a slight over-tilt and under-tilt of ±0.5° (Figure S3). During the final 2 kV-polishing, we used an over and under tilt of ±10° to produce a thin, low-damage region extending 4–7 µm from the front of the lamella with minimal change in the ion penetration depth (Fig. S4a, Fig. 1a,d,e). The high milling angle was essential because milling with a ±4°tilt only removed material within the front ~1.5 µm of the lamella, leaving the remainder of the lamella with damage from the prior 8 kV step (Fig. S4b,c) (Fig. 1d). A more detailed description of this method can be found in the methods and (Supplementary Fig. S3).

**Figure 1.**
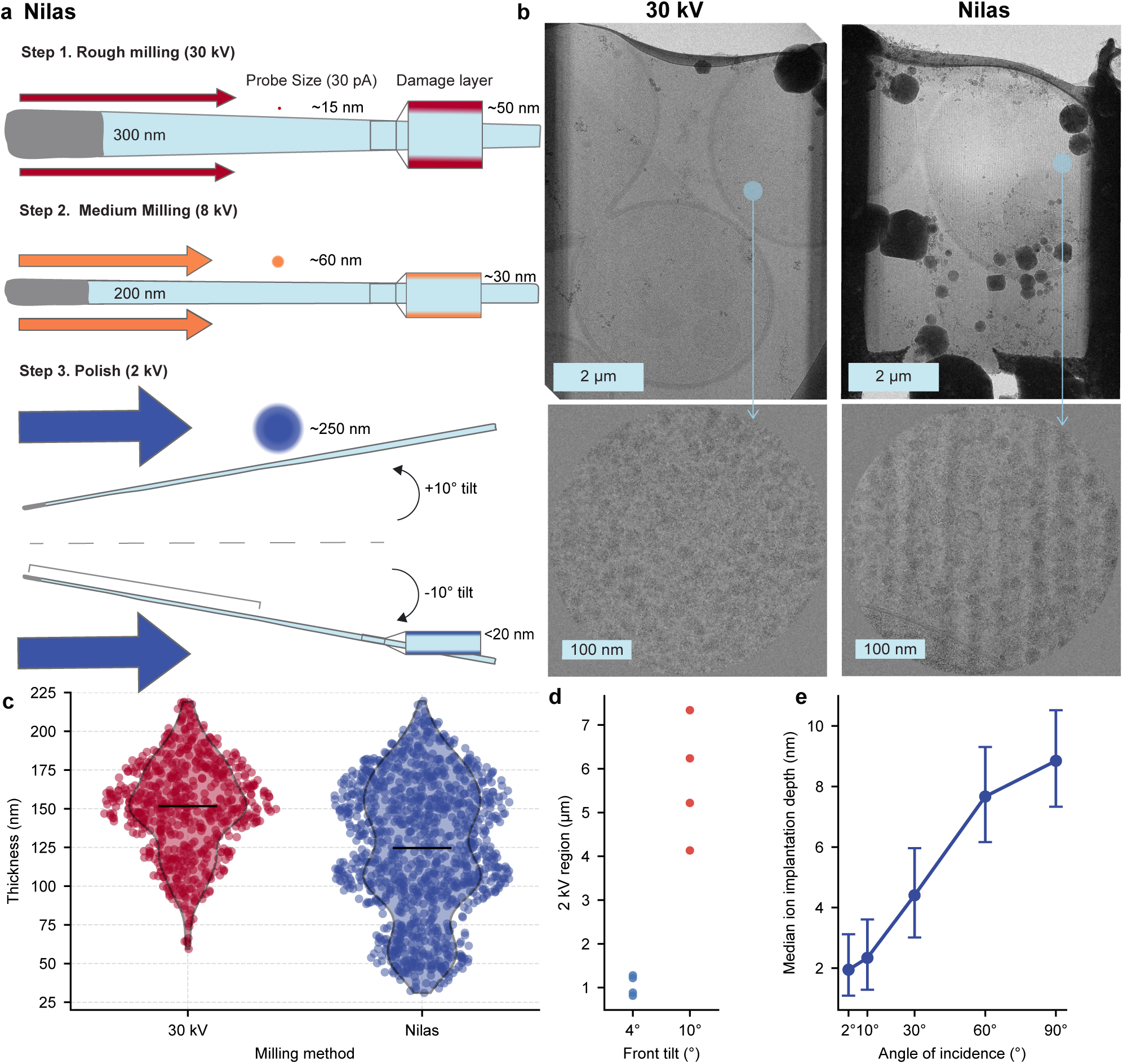
Nilas results in thin, 2 kV polished lamellae over large areas. (**a**) Diagram of Nilas milling method with steps at 30 kV, 8 kV and 2 kV. Probe size for gallium at the typical polishing current of 30 pA depicted. The depth of the FIB-damage layer that is caused by milling with each accelerating voltage is shown. (**b**) Example TEM images of lamellae milled with standard 30 kV polishing and Nilas. Cytoplasmic high-magnification TEM images from each lamellae. (**c**) The distribution of CTFFIND5 estimated thicknesses for images collected on Nilas and 30 kV lamellae. (**d**) Length of the 2 kV polished lamella region when polished at ± 4 and ± 10 degrees. (**e**) SRIM simulated median depth of ion implantation for a gallium 2 kV ion beam in a frozen yeast cell. Error bars indicate the inter-quartile range (IQR).

The sputter rate, and consequently milling speed, of gallium at 2 kV is ~5 times lower than gallium at 30 kV ^27^. The voltage step-down approach enables Nilas to remove the damage layer while maximizing efficiency. The final 2 kV polish removes the ~30 nm of damage from the 8 kV milling step in *<*30 seconds per side (Fig. 1a).

We named this protocol Nilas, an oceanic term for thin sea ice, because using this method we were able to consistently generate thin lamellae with sub 50 nm regions, and the entire area occupied by a yeast cell at sub-100 nm thickness, which was not observed in any 30 kV-milled lamellae (Fig. 1b,c). Images of thin lamellae regions show a visible increase in contrast (Fig. 1b).

### Nilas reduces FIB-milling damage

We compared the FIB-damage in Nilas-milled lamellae to 30 kV-milled lamellae on the same grids milled in the same session using the LSU 2DTM damage assay as described above (Fig. 2a) (Supplementary Fig. S5a). LSUs identified in Nilas-milled lamellae have higher 2DTM z-scores relative to 30 kV-milled lamellae of similar thickness and are further increased in thin sections (Fig. 2b-d). We observed broader plateaus of undamaged LSUs and significantly less damage in Nilas-milled lamellae relative to 30 kV-milled lamellae at depths up to 50 nm (*p*<0.05, t-test)(Fig. 2b,e). Consistently, we observed a higher LSU density towards the lamella surface and for lamellae of a comparable thickness (Supplementary Fig. S5b,c). We integrated the effects of FIB-damage and multiple and inelastic electron scattering ^9^, using a previous estimate of the 2DTM decay constant *λ*_SNR_ ^28^. Consistent with our prior work ^9^, we predict that recovery of signal from 30 kV-milled lamellae is maximal in sections of ~ 90 nm (Fig. 2f). In contrast, Nilas-milled lamellae are closer to the physical signal limit imposed by electron scattering across the full range of lamella thicknesses and have a maximal signal recovery at ~40 nm (Fig. 2f). These predictions are consistent with the observed maxima of the mean 2DTM z-scores per image and an increase in maximal signal in the Nilas-milled images of ~20% (Fig. 2g). We no longer observe a difference between the mean 2DTM z-score of LSUs identified in Nilas-and 30 kV-milled lamellae in images ≥150 nm. A smaller proportion of identified LSUs will be damaged in thicker samples and fewer damaged ribosomes will be detected due to the overall decrease in signal which may explain this observation.

**Figure 2.**
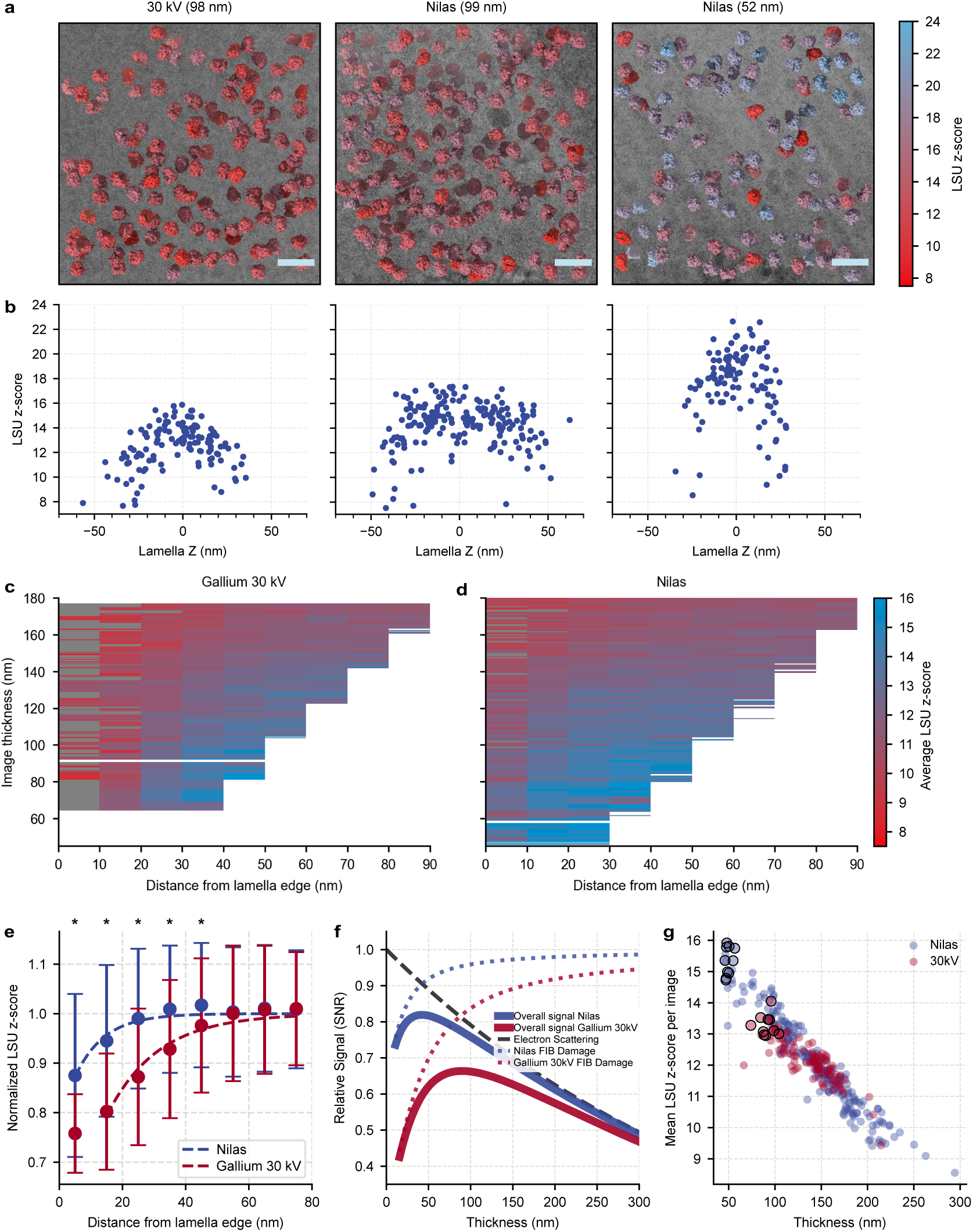
Nilas dramatically reduces FIB-milling damage. (**a**) Example images from 100 nm thick sections of 30 kV-and Nilas-milled lamellae overlaid with volumes at the locations and orientations of significant LSUs detected by 2DTM and colored by 2DTM z-score. Right: an example image of a thin (50 nm) section enabled by Nilas with the same annotation. Scale bar: 50 nm. (**b**) Scatterplot showing the 2DTM z-scores of significant LSU detections in the above images plotted as a function of their lamella z-coordinate as determined by 2DTM. (**c,d**) Heat maps of the mean 2DTM z-score from 10 nm bins of lamellae at indicated thicknesses for both milling conditions. Grey indicates fewer than three ribosomes detected in the bin. (**e**) Scatterplot of the mean 2DTM z-scores of LSUs binned by distance to lamella edge normalized to undamaged bins in the center of images in Nilas-milled (blue) or 30 kV-milled (red) lamellae. Curves represent the fits of the exponential decay function *y* = 1 *− Y*_0_*e^−d/k^*. Fit parameters Nilas: *Y*_0_ = 0.22*, k* = 9.3, *R*^2^ = 0.94; gallium 30 kV: *Y*_0_ = 0.52*, k* = 16.1, *R*^2^ = 0.96. Asterisks indicate a significant (*p <* 0.05, t-test) difference between the two conditions at the indicated depth bin. Error bars indicate ± a standard deviation. (**f**) Plot showing the predicted SNR as a function of lamella thickness integrating FIB-damage and electron scattering for 30 kV-and Nilas-milled lamellae. (**g**) Scatterplot showing the mean LSU 2DTM z-score per image as a function of measured thickness and milling condition. The ten circled data points represent the 10 best images from each condition used in subsequent analysis.

We only observed evidence of damage within 20 nm of the surface in Nilas-milled lamellae (Fig. 2b). The location of the 2DTM peak corresponds to the center of mass of the LSU template. Therefore any particle found within a ribosome radius (~15 nm) (Supplementary Fig. S5a) from the surface is partially ablated, resulting in decreased 2DTM z-scores, independent of sub-surface FIB damage. A simulated idealized damage curve that accounts for partially ablated ribosomes alone resembles the Nilas damage curve, suggesting that our measurement is limited by the size of the LSU (Supplementary Fig. S5d). Across multiple images, Nilas-milled lamellae have more LSUs detected within 10 nm of the lamella surface and in several cases the highest mean binned z-score was within 20 nm of the lamella surface (Fig. 2c,d). Therefore we expect that we are overestimating the damage in Nilas-milled lamellae. For this reason, we predict to see further improvement in SNR for smaller complexes in lamellae thinner than the predicted optimal of *∼*40 nm.

### Nilas improves the resolution and interpretation of *in situ* 3D reconstructions

We next tested whether the increased SNR in images of thin sections of Nilas-milled lamellae improves the downstream analysis of particles within these lamellae. The most common application for *in situ* structural biology is the determination of high resolution structures by averaging identified particles. We selected images representing the highest LSU 2DTM z-scores from each condition. We identified 3,113 and 2,748 significant detections in 40 images from Nilas-and 30 kV-milled lamellae respectively using a 1 MDa fragment of the LSU as a template for 2DTM (Supplementary Fig. S6a,b). We used baited reconstruction ^29^ to generate 3D reconstructions using the locations and orientations from 2DTM of 2,748 particles from each condition. Both the global (4.94 Å relative to 5.93 Å) and local resolution of the 3D reconstruction from Nilas-milled lamellae was higher for the same number of particles relative to 30 kV-milled lamellae (Supplementary Fig. S6 c,d) (Fig. 3a). Consistently, the Rosenthal-Henderson B-factor ^30^ was lower for particles from Nilas-milled lamellae (51 Å^2^) relative to 30 kV-milled lamellae (81 Å^2^) (Fig. 3b) which we expect is a result of the reduction in both damage and sample thickness (Supplementary Fig. S6b). To control for template bias, we compared the global and local resolution of the LSU in regions omitted from the 1 MDa truncated template (Supplementary Fig. S6a). We observed an improved global (4.31 Å relative to 4.60 Å) and local resolution throughout the omitted region for the reconstruction from Nilas-milled lamellae relative to the reconstruction from 30 kV-milled lamellae (*p <* 0.05, block bootstrap) (Fig. 3c,d) (Supplementary Fig. S6e,f).

**Figure 3.**
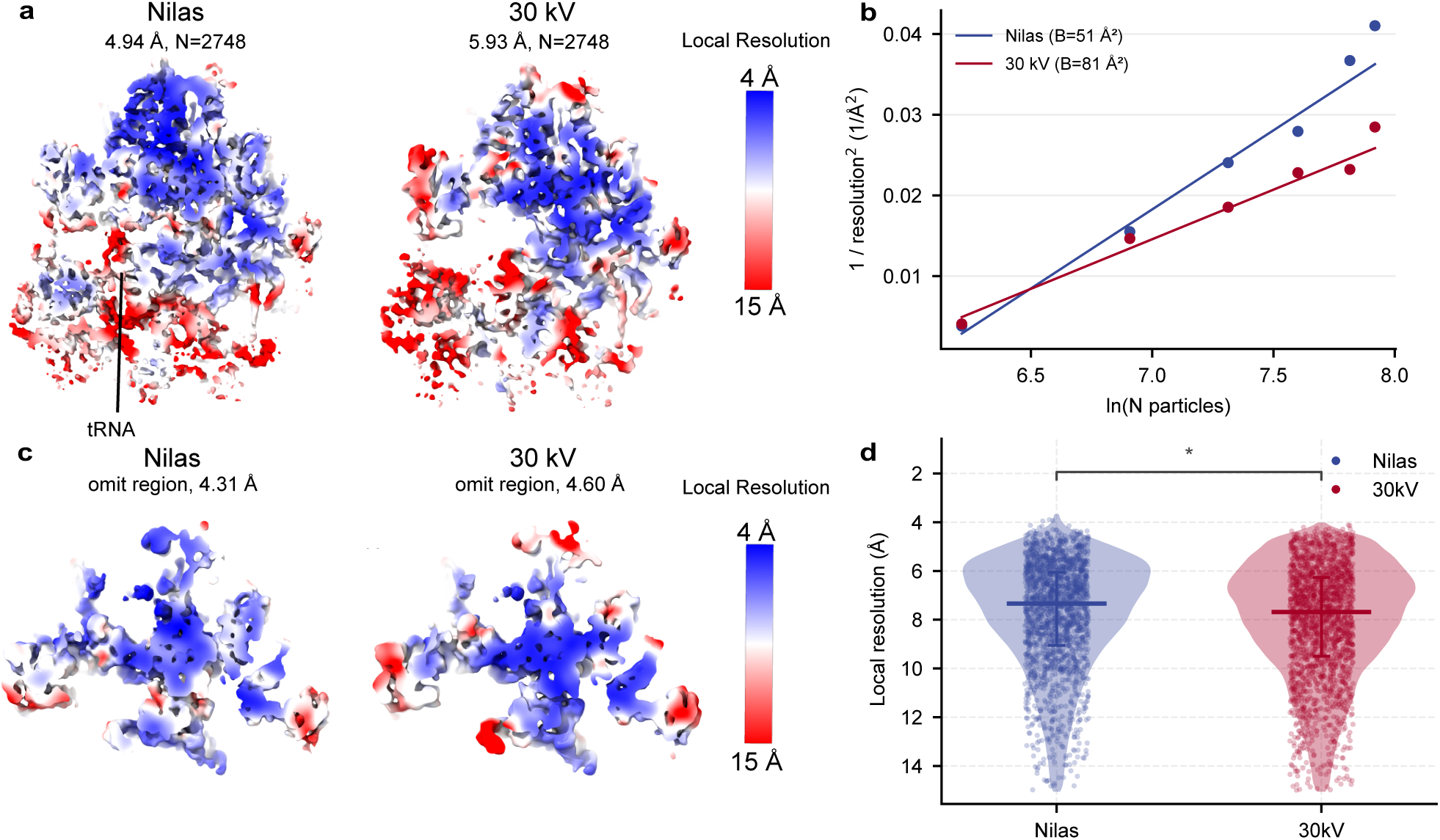
Nilas improves the resolution of 3D reconstructions. (**a**) 3D reconstructions of Nilas and 30 kV particle sets, colored by local resolution. Label shows a density consistent with a tRNA present in the Nilas reconstruction. (**b**) Rosenthal-Henderson B-factor plot of the resolutions of reconstructions generated from random particle subsets. (**c**) Visualization of the region of the reconstruction omitted from the template and colored by the local resolution for each condition. Resolution measured by masked FSC of omitted region shown. (**d**) Local resolution plotted per voxel for the omitted LSU region of Nilas and 30 kV reconstructions shown in (c). Median and IQR annotated on plot. Values were filtered to better than 15 Å. Asterisk indicates *p <* 0.05 (see methods).

We observed density consistent with a tRNA, not included in the template, in the reconstruction from Nilas-milled lamellae, but not in the reconstruction from 30 kV-milled lamellae. This shows that the improved SNR in Nilas-milled lamellae directly translates to improved characterization of biologically relevant structural features (Fig. 3a).

### Nilas improves the recovery of ribosomal subunits and extends the size limit for detection with 2DTM

The body of the small ribosomal subunit (SSU; PDB: 6Q8Y) ^25^, while relatively large at ~600 kDa, has been difficult to identify with 2DTM relative to the ~1.8 MDa LSU^16,17,31^. We expect LSUs and SSUs to be roughly stoichiometric. However, in a previous study with cells in similar conditions, we identified 10x fewer SSUs than LSUs in independent searches ^31^. In thin sections of Nilas-milled lamellae, we identify SSU bodies up to ~65% of the number of detected LSUs without incorporating prior information about the LSU. We also noted that, even for the same thickness, Nilas-milled lamellae tend to recover more SSUs relative to 30 kV-milled lamellae (Fig. 4a,b).

**Figure 4.**
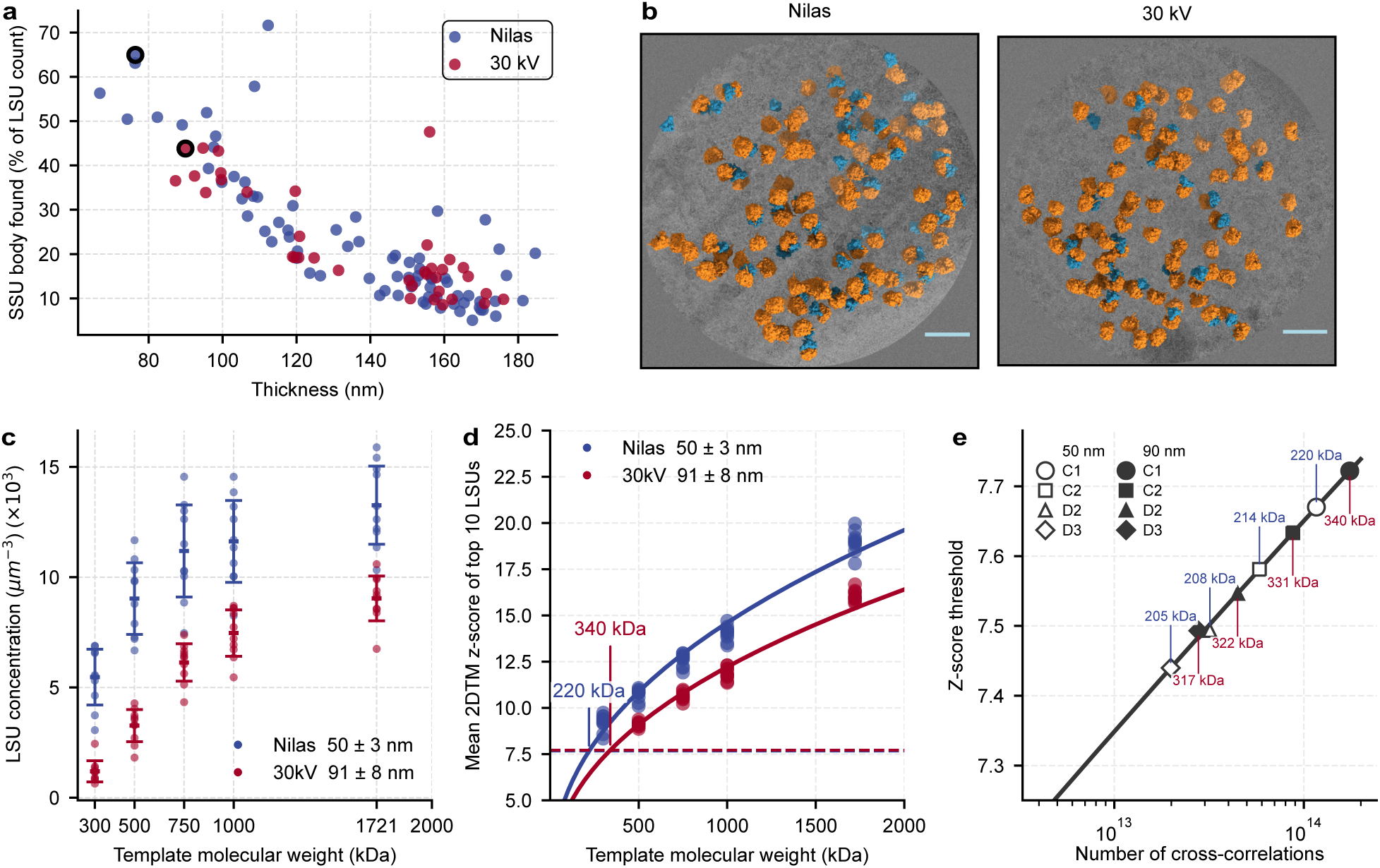
Nilas improves the recovery of ribosomal subunits. (**a**) Scatterplot showing the SSU detection rate expressed as a percentage of LSUs detected in the same image as a function of thickness. Outlined points are selected images displayed in (Fig. 4b) (**b)** Representative 30 kV and Nilas images showing LSUs (orange) and SSUs (blue) identified by 2DTM. Scale bar: 50 nm. (**c**) LSU concentration with truncated ribosome templates for the ten best cytoplasmic images per condition. (**d**) Scatterplot showing the mean 2DTM z-score of the top 10 peaks in each image as a function of the molecular mass of the truncated LSU template. Dashed line indicates the 2DTM z-score threshold for one false positive per image. Data were fit to *Z* = *a M^k^* with *k* shared between conditions, yielding *k* = 0.43, *a* = 0.77 for Nilas lamellae and *a* = 0.65 for 30 kV lamellae (*R*^2^ = 0.97 for both). (**e**) The 2DTM z-score threshold as a function of the number of cross-correlations in the 2DTM search. Points along the line indicate the significance thresholds allowing a single false positive per image for common searches and symmetries. Thicker images require the searching of more defocus planes. Images of 100 nm sections require sampling 9x 200 Å defocus planes, whereas 50 nm regions require sampling only 6x 200 Å defocus planes. Colored molecular weight labels indicate the predicted minimal detectable molecular weight for Nilas (blue) and 30 kV-milled (red) images.

To compare the recovery of smaller complexes in Nilas-milled lamellae we selected ten images representing the maximal information recovery for each milling type, (outlined in Fig. 2g) and used 2DTM with a set of truncated LSU templates (300 kDa–1 MDa). We observed an increased density of LSUs in images of Nilas-milled lamellae at all molecular weights, from ~1.45x with the full length template to ~4.6x with the 300 kDa template (Fig. 4c). The 500 kDa template in Nilas-milled lamellae recovered a similar density of ribosomes as with the full template in 30 kV images (Fig. 3c). We expect that the observed increased density results from both the reduced damage and improved SNR in thinner sections from Nilas-milled lamellae. Because thinner sections will have a higher proportion of damaged particles, we expect the improvement from thin Nilas-milled lamellae to be greater for smaller particles. To predict the minimal detectable molecular weight, we examined the mean 2DTM z-score of the top 10 significant detections in each image for each truncated template as a function of molecular mass ^16^. The mean 2DTM z-score was consistently higher for Nilas-milled lamellae across molecular weights (Fig. 4d). The relationship between 2DTM z-score (*Z*) and molecular weight (*M*, in kDa) was modeled by an exponential function (Fig. 4a,b). The resulting fit was close to the expected SNR scaling with the square-root of the molecular mass ^7,8^, the small deviation (exponent = 0.43) may result from the 2DTM projection normalization slightly deflating the z-score of larger templates. The fitted model predicted a minimal detectable molecular mass of 220 kDa with Nilas and 340 kDa for 30 kV polished lamellae (Fig. 4d). The threshold used to call significant detections is based on the probability of a false alarm from a Gaussian noise distribution and depends on the number of cross-correlations performed ^7,31^. Thinner samples require fewer defocus searches relative to thick samples and template symmetry permits sampling fewer orientations reducing the z-score threshold and predicted minimal detectable molecular mass (Fig. 4e).

### Thin, undamaged lamellae improve the recovery of non-ribosomal complexes *in situ*

We observed improved detection of ribosomal subunits with 2DTM in images of thin, Nilas-milled lamellae. However, ribosomes are abundant and largely RNA, which has improved contrast in cryo-EM relative to proteins and is more resistant to radiation damage. To test whether Nilas also improves the recovery of other macromolecules we used 2DTM to localize complexes representing differing abundance and over an order of magnitude in molecular mass. To minimize false positives when detecting lower abundance complexes, we used a significance threshold that permitted a single false positive over 10 images searched for all complexes.

We identified 7 instances of fatty acid synthase in 6/10 cytoplasmic images (FAS, 2.6 MDa, PDB:2UV8) ^32^ from Nilas-milled lamellae and 15 in 8/10 cytoplasmic images from 30 kV-milled lamellae (Fig. 5a-d). If evenly distributed in the cytoplasm, we estimate there should be ~1 FAS per image, suggesting we are finding all or nearly all molecules present for both searches ^33,34^ BNID:110680. The average and maximum 2DTM z-scores were ~two standard deviations higher for the detections in Nilas-milled lamellae (Fig. 5e). We conclude that the improvement in SNR in Nilas-milled lamellae is not isolated to ribosomes, and extended our search to other complexes.

**Figure 5.**
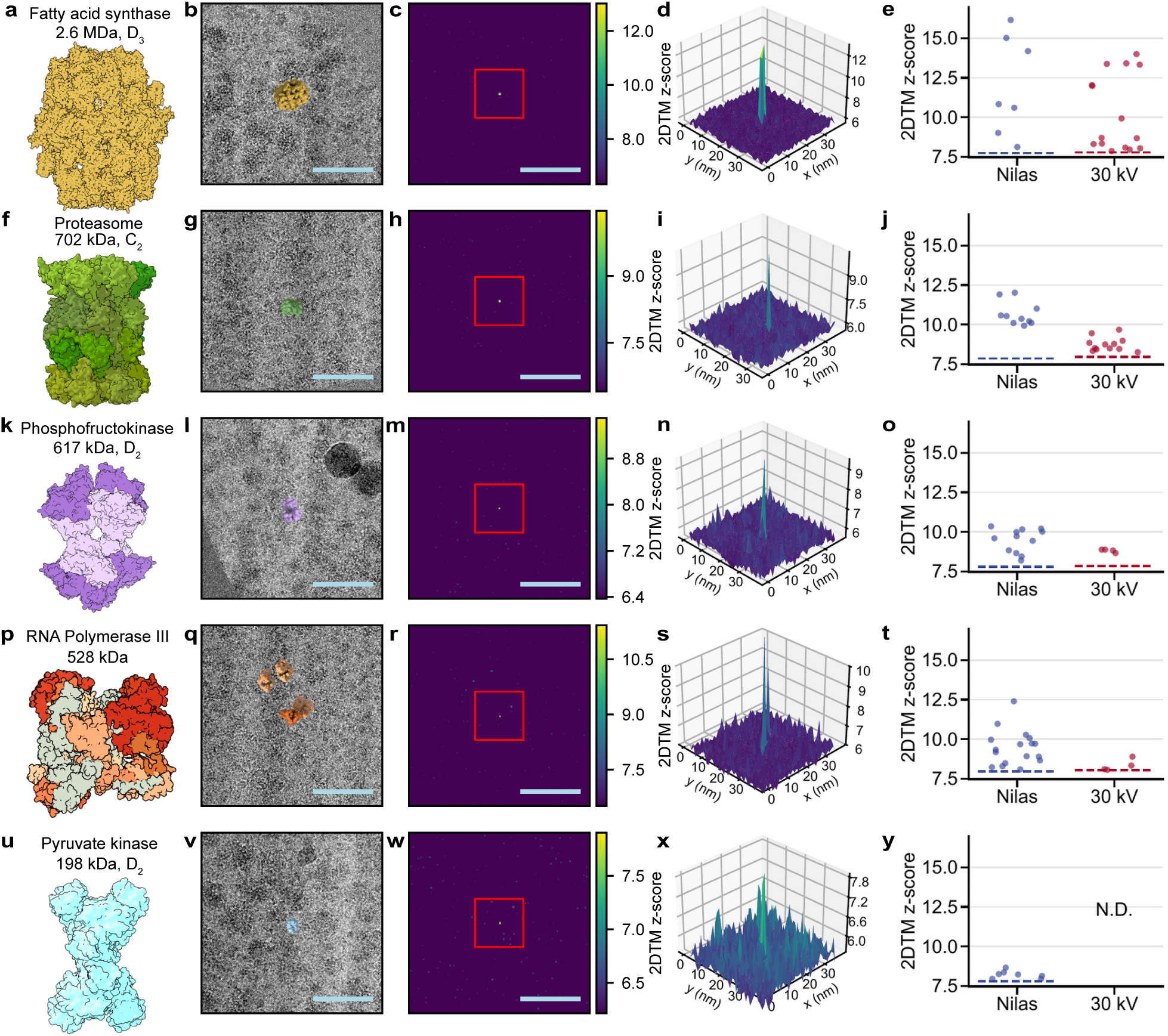
Nilas improves the recovery of diverse complexes with 2DTM. (**a)** Molecular model of fatty acid synthase (FAS; PDB: 2UV8) used as a 2DTM template. (**b)** Overlay of FAS template at a significant location and orientation identified by 2DTM on a cropped image region. Scale bar: 50 nm. (**c)** Scaled maximum intensity projection (MIP) representing per pixel 2DTM z-score of the cropped region from (b). (**d)** 3D visualization of the cropped region from red box in (c), centered on the highest peak. (**e**) Significant 2DTM z-scores for FAS in ten cytoplasmic Nilas-and 30 kV-milled lamella images. Dashed line represents the 2DTM z-score threshold for 1 false positive per 10 images. (**f)** Molecular model of the 20S proteasome (PDB: 1RYP) used as a 2DTM template. (**g)** Overlay of the proteasome template at a significant location and orientation identified by 2DTM on a cropped image region. Scale bar: 50 nm. (**h)** Same as (c) but for (g). (**i)** Same as (d) but for (h). (**j)** Significant 2DTM z-scores for indicated targets in ten nuclear Nilas and 30 kV images. Dashed line represents the 2DTM z-score threshold for 1 false positive per 10 images. (**k**) Molecular model of phosphofructokinase (PDB: 3O8O) used as a 2DTM template. (**l**) Overlay of phosphofructokinase template at a significant location and orientation identified by 2DTM on a cropped image region. Scale bar: 50 nm. (**m**) Same as (c) but for (l). (**n**) Same as (d) but for (m). (**o**) Significant 2DTM z-scores for indicated targets in ten cytoplasmic Nilas-and 30 kV-milled lamella images. Dashed line represents the 2DTM z-score threshold for 1 false positive per 10 images. (**p**) Molecular model of RNA polymerase III (PDB: 8BWS) used as a 2DTM template. (**q**) Overlay of RNA polymerase III template at a significant location and orientation identified by 2DTM on a cropped image region. Scale bar: 50 nm. (**r**) Same as (b) but for (q). (**s**) Same as (c) but for (r). (**t**) Significant 2DTM z-scores for indicated targets in ten nuclear Nilas-and 30 kV-milled lamella images. Dashed line represents the 2DTM z-score threshold for 1 false positive per 10 images. (**u**) Molecular model of pyruvate kinase (PDB: 1A3X) used as a 2DTM template. (**v**) Overlay of pyruvate kinase template at a significant location and orientation identified by 2DTM on a cropped image region. Scale bar: 50 nm. (**w**) Same as (c) but for (v). (**x**) Same as (d) but for (w). (**y**) Significant 2DTM z-scores for indicated targets in ten cytoplasmic Nilas-and 30 kV-milled lamella images. Dashed line represents the 2DTM z-score threshold for 1 false positive per 10 images.

We identified 10 instances of the 20S proteasome (702 kDA) PDB: 1RYP) ^35^ from 6/10 nuclear images from Nilas-milled lamellae and 12 proteasomes in 7/10 nuclear images from 30 kV lamellae (Fig. 5f-j). Proteasomes identified in Nilas-milled lamellae were consistently ~2 standard deviations higher and exceeded the significance threshold by almost 3 standard deviations (Fig. 5f-j). Proteosomes were more frequent in images close to the nuclear periphery. One image from a Nilas-milled lamella had four significant detections, three of which are adjacent to a visible nuclear pore, consistent with prior observations in *Chlamydomonas reinhardtii* (Supplementary Fig. S8a) ^36^.

Phosphofructokinase (617 kDa; PDB: 3O8O) ^37^ is the rate limiting enzyme in glycolysis, and is allosterically regulated to control flux through glycolysis. Therefore, understanding its structural and oligmetric state *in situ* is important to understanding its function and regulation. We identified three times as many significant detections in Nilas-milled lamellae relative to 30 kV-milled lamellae (Fig. 5k-n), corresponding to a ~5.5x higher density (Fig. 5o). While the number of overall detections is low, the 2DTM z-scores allow us to establish that they are unlikely to be false positives (Supplementary Fig. S7). We find ~50% of the expected number of PFK molecules in Nilas images and 10% in 30 kV images ^33,34^, lower than the recall of 500 kDa truncated ribosomes (Fig. 3f). This could be due to the aspherical nature of PFK which makes the z-score a less reliable metric ^38^, the absence of RNA in the template, structural differences between the PFK template and targets or inhomogenegous distribution of PFK in the cytoplasm.

RNA polymerases are ~10x less abundant than ribosomes and localize to the nucleus where they have been difficult to identify using cryo-ET or 2DTM. The ~3-fold improved recovery of 500 kDa LSU fragments in Nilas-milled lamellae predicts that detection of RNA polymerase III should be possible. Indeed, we recover 17 instances of RNA polymerase III (528 kDa, PDB: 8BWS) ^39^ in 10 cytoplasmic images of Nilas-milled lamellae compared to 4 instances in 4/10 images from 30 kV-milled lamellae (Fig. 5p-t). This exceeds the expected number if the polymerase was uniformly distributed in the nucleoplasm ^33,34^. Mapping the template to the locations and orientations identified with template matching allowed us to observe cases where the DNA binding grooves of RNA polymerase III molecules are aligned (Supplementary Fig. S8b), consistent with transcribing the same strand of DNA. We did not find any significant detections in three images of the cytoplasm, where we do not expect to find RNA polymerase III. Moreover, in several nuclear images we observed dark spots with a periodicity of ~3.5 nm, consistent with one full turn of a DNA duplex, based on the structure of B-DNA (PDB: 4BNA) ^40^ (Supplementary Fig. S8 b,c). These patterns were often surrounding and extending before or after the polymerase, which we hypothesized is the direct visualization DNA facilitated by the higher contrast in images of thin sections. In one case a super-helical pattern is observed which could indicate super-coiled DNA (Supplementary Fig. S8 c). However, due to the thin region visualized, we cannot unambiguously trace a DNA strand in three-dimensions.

More abundant particles will have an increased probability of detection above the significance threshold, predicting possible detection of some small, highly abundant complexes. Pyruvate kinase is a glycolytic enzyme which catalyzes the final irreversible step of glycolysis, and is one of the most abundant proteins in the cell. The pyruvate kinase tetramer at 198 kDa is just below the minimal detectable molecular mass based on estimates from the LSU fragments. Despite its small size, we recovered ten significant detections (0.2% of the number of predicted molecules ^33,34^) of pyruvate kinase (198 kDa, D2 symmetry, PDB: 1A3X) ^41^ in Nilas-milled lamellae (Fig. 5u-y). We did not recover any significant detections in images of 30 kV-milled lamellae. These detections are unlikely to be false positives based on the observed 2DTM z-scores (Supplementary Fig. S7) and the 2DTM z-score threshold of 1 false positive per 10 images. Moreover, we did not find any significant detections in three images of the nucleus, where we do not expect to find pyruvate kinase. This highlights the importance of considering target abundance in addition to molecular mass.

## Discussion

Standard protocols for FIB milling introduce damage that destroys particles and eliminates the benefit of thinning below 100 nm. FIB damage limits annotation of the proteome and the determination of high resolution structures *in situ* to large and abundant complexes. Here, we describe Nilas, a FIB-milling protocol optimized for visual proteomics which improves the detection of macromolecules through the generation of thin, minimally damaged lamellae. We demonstrate that Nilas significantly reduces FIB-milling damage throughout the lamella and dramatically improves the recovery of signal from lamellae <150 nm relative to standard milling protocols. Nilas-milled lamellae produce higher resolution reconstructions and improve the interpretation of biological features with fewer particles. With Nilas, the recovery of both ribosomal subunits and important, non-ribosomal complexes with 2DTM is improved. Notably, Nilas-milled lamellae enabled us to map RNA polymerase III in the yeast nucleus with 2DTM which has not previously been identified. While we used 2DTM in untilted micrographs to maximize sensitivity of the damage assessment, our measurements apply directly to cryo-ET. By developing a FIB-milling protocol optimized for visual proteomics, we expand the impact of this technology, bringing us closer to annotation of the structured proteome.

### Prospects for further improving lamella quality with FIB-milling

While Nilas minimizes damage compared to standard milling strategies, thinning by collisions with ions of any energy will result in some damage. Based on predictions from stopping range of ions in matter (SRIM) simulations, we expect that further reducing the accelerating voltage below 2 kV will have diminishing returns on reducing the effective damage layer. Reducing the accelerating voltage to 0.5 kV, for example, would only be expected to reduce the median ion penetration depth by about 10 Å, below the size of most proteins and below the accuracy of defocus estimation in 2DTM and z-resolution in tomography. Additionally, at lower voltages there is an increased disparity in sputter yields for materials of different densities, which results in preferential milling of less dense areas. Therefore, further reducing damage would require new milling strategies. Possibilities include the integration of low-energy (100–500 eV) argon polishing ^42^, gas cluster ion beam irradiation (GCIB) ^43^ or plasma, as has previously been proposed ^44^. At a few eV individual ion energies, GCIB offers broad, shallow energy deposition, that due to different cluster masses is well suited to producing smooth, minimally damaged lamellae ^43^.

The final 2 kV milling step in Nilas introduces fine curtaining with an approximate 20 nm frequency (Fig. 1b). The curtaining does not affect identification with 2DTM (Fig. 2a, b). However, curtaining is a strong low-resolution feature that can affect alignment quality of tomograms and will be accentuated by the improved contrast with the use of a phase plate. Therefore, a smooth and thin sample is ideal to permit further imaging applications with Nilas-milled lamellae. Curtaining at low voltages can result from non-uniform dose across the lamella as dose calculations are tuned to 30 kV probe size and beam shape. Additionally, at low voltages, increased aberrations distort the beam away from a Gaussian profile, producing beam tails that do not mill as expected even when dose is correctly tuned ^20^. To mitigate these effects, Nilas utilizes an increased thickness of the GIS layer, which homogenizes the surface the beam first encounters, an increased beam overlap (85%) to account for the beam profile and a multi-directional scanning pattern to further reduce directional milling artifacts (Supplemental Fig. S3). In the future, rock milling, in which samples are milled from multiple directions to minimize the curtaining artifact, could offer further improvement ^20,45^.

Thin lamellae milled with Nilas are more fragile and susceptible to breakage, therefore there is a tradeoff between throughput and lamella thickness. Of 22 lamellae attempted, 18 survived 8 kV polishing and after 2 kV polishing, four were unsuable, and of the 14 usable lamellae, three were intact and of high quality, five contained holes or cracks but had imageable areas, and six appeared high quality and intact on the FIB (confirmed by SEM) but broke during transfer between the FIB and TEM (Supplementary Fig. S9a-d). These lamellae frequently broke at the boundary between the cell wall and cell membrane (Supplementary Fig. S9)a. Cells without the cell wall-cell membrane boundary, such as mammalian cells, may show a higher recovery. Loss of intact lamellae during transfer might also be mitigated by improved handling, for example by transferring directly from the FIB-SEM to the TEM, as is the case with the Arctis cassette transferred directly to the Krios (Thermo Fisher) and the cryolamellar to the cryoARM300 (JEOL). In this experiment our goal was to generate very thin lamellae, which contributed to the lower yield.

We observed increased inter-frame motion in some images of *<*90 nm, and no difference in thicker regions, relative to 30 kV-milled lamellae (Supplementary Fig. S9e). Lamellae flexing is a result of thermal and mechanical stress which can be exaggerated in thinner, less mechanically stable lamellae ^46^. Altering lamella geometry could make lamellae more stable in general ^47^ or increase the stability of thin regions in particular ^48^ but will reduce the very thin area.

While currently manual, automation of Nilas with standard instrumentation could increase throughput. Public software projects such as fibsemOS could be adapted to perform Nilas ^49,50^. Proprietary software such as AutoTEM (Thermo Fisher) would require changes from the manufacturers to enable automated milling.

### Comparison with other low energy methods

A recent preprint describes a low energy polishing (LEP) method that uses 8 kV gallium polishing to improve lamella quality ^22^. Similar to the first two steps of Nilas, they use an increased deposition of organo-platinum with a GIS, and a slight over and under tilt for the 8 kV milling step, which we find helps protect the GIS layer. We observed that 8 kV polished lamellae showed decreased damage relative to 30 kV polished lamellae. However, polishing with 2 kV results in a significant further reduction in damage (Supplemental Fig. S4a) and can produce thinner lamellae. We observed a similarly high success rate after the 8 kV polishing step (~85%) and minimal curtaining (Supplemental Fig. S4,S9e). Where throughput is a higher priority than maximum signal, or the application is sensitive to curtaining artifacts, an 8 kV polish is the optimal protocol. In order to mill lamellae that have maximally thin regions with minimal damage, the 2 kV polish offers an advantage.

Although Nilas and LEP use gallium LMIS as the FIB, Nilas could be applied with plasma FIBs using protocols that account for differences in beam profiles as has been done for the transmission electron microscopy (TEM) preparation of polycrystalline aluminum thin films on silicon wafer ^24^. We selected gallium because it is more focused when using low currents at 2 kV relative to xenon and is more broadly available to users. However, the minimal requirement for Nilas is to be able to identify the lamella site, and our initial testing suggests standard 8 kV and tilted 2 kV polishing with xenon is workable. New plasma FIB columns, with decreased spot sizes for xenon at low currents will be advantageous for this process ^51^.

### Capturing cellular context

A downside of FIB-milling thin sections is the dwindling cellular context within a thin region. Reducing lamella thickness to 50 nm or less further reduces the volume imaged. In this work, we have demonstrated that the improved SNR in thin samples enables the detection of important complexes such as ribosomal subunits, phosphofructokinase, the proteasome and polymerase III. The optimal tradeoff between lamella volume and detection will depend on the biological question at hand. If thicker lamellae are important for the biological question, minimizing damage with Nilas will still improve data quality.

Context in all dimensions must be considered. Increasing cellular context in x,y with the use of montage imaging strategies like DeCo-LACE^52^ and montage tomography ^53–56^, can cover a ~600-fold greater area in x,y with beam image shift, relative to the 4-fold difference between a 200 nm lamella and a 50 nm lamella, without the resulting loss in signal from a thicker sample. The ability to capture more of the proteome, over a larger area, provides greater potential for recovering cellular context relative to slightly thicker lamellae.

Our work has shown the potential gain from the improved SNR and visual contrast in very thin cellular samples. Ideally, we would not have to tradeoff SNR and cellular context by generating serial thin sections. To date, the only method with the potential to generate serial thin samples is CEMOVIS^5^. Further developments are required to make CEMOVIS an effective strategy to visualize the proteome.

### Limitations in measuring FIB-milling damage

The ability to accurately quantify FIB-milling damage depends on the size of the ruler used to measure it, since larger macromolecules will be partially ablated by surface removal even in the absence of further subsurface damage. In 2DTM, the position of the particle is defined by its center of mass and is limited by the radius of the particle which is 15 nm for the LSU. In thin images, the LSUs 10–20 nm from the lamella surface of Nilas-milled lamellae sometimes contain the highest 2DTM z-scores in an image, consistent with damage not extending beyond 10 nm (Fig. 3d). Therefore, for particles smaller than the LSU, we are likely overestimating the damage remaining in Nilas-milled lamella. We observe a clear and consistent reduction in the damage layer in Nilas-milled lamellae relative to 30 kV-milled lamellae, indicating that in 30 kV-milled lamellae damage extends beyond the surface and is not only the result of ribosomes being cut during milling. A more accurate quantification of subsurface damage would require an abundant, <5nm diameter particle that can be easily detected. A small particle will not be detected at rates similar to the ribosome and would have more uncertainty in defocus estimation, thereby reducing the accuracy of the metric with 2DTM.

### Detection of more of the visual proteome

With Nilas it is possible to generate biological lamellae of a thickness comparable to single particle samples. With traditional 30 kV polishing, lamellae are typically 150 nm–250 nm thick. The increase in SNR by reducing the lamella thickness from 50 nm from 200 nm is ~1.5 fold, decreasing the minimal detectable molecular mass by a factor of ~2.25.

Improved visual contrast in thinner lamellae may also play a role in detection. The whitening filter currently used with 2DTM suppresses low-resolution features and may not prove optimal for particle detection in thin lamellae. Future improvements could be seen through customizing filters and search strategies to optimize detection, or by balancing low and high-resolution features with metrics like the p-value ^38^. Effective hybrid collection schemes combining 2D imaging with subsequent tomograms could also improve annotation ^17,57,58^.

The ability to generate thin lamellae also has implications for the utility of the laser phase plate (LPP) ^59,60^ for detection of molecules in cellular sections. Imaging in focus with a 90 degree phase shift could give up to a 2-fold improvement in contrast by removing CTF zeros ^7,61^, further increasing SNR and improving the minimal detectable molecular mass. However, the benefit drops quickly further from focus such that if a 200 nm section is aligned with its center in focus, the edges would no longer gain the same benefit of in focus imaging. Nilas makes in-focus imaging of lamellae with a laser phase plate feasible.

Further gains could be achieved by combining Nilas and LPP imaging with other recent hardware developments. Reducing radiation damage with liquid helium cooling could further reduce the minimal detectable molecular mass by 1.5 fold ^62^. Correcting chromatic aberration (Cc) in thick samples could allow for recovery of signal from inelastically scattered electrons ^63,64^. In the sub-50 nm sections we observed in Nilas-milled lamellae, lowering the TEM beam energy to 100-200 keV in place of a 300 keV electron beam could further reduce the predicted minimal detectable molecular mass ^65^.

In combination with algorithmic improvements it is within reasonable possibility that much more of the structured proteome could be annotated with 2DTM. Detecting particles using the cross-correlation of high-resolution features enforces a reliance on consistency between the template and target molecule for detection. This precludes detection of more conformationally dynamic complexes. Reducing the ordered molecular mass required for detection will allow for identification of some dynamic complexes by using a smaller, more locally ordered region in a complex as a template. This could shift *in situ* cryo-EM from a method for determining the structure of molecular complexes in their native cellular environment to one capable of producing molecular atlases of the cell.

## Methods

### 4.1 Cell culture and grid preparation

*Saccharomyces cerevisiae* strain W303 haploid (NucLoc) or BY4741 colonies were grown to mid-log phase (0.3–0.8) at 30°C, 200 RPM. Yeast was diluted to 15,000 cells ml*^−^*^1^ and 5 µl was applied to a Quantifoil R 2/2 SiO2 200 mesh Cu grid or an Ultrafoil R 2/2 200 mesh Au grid. Grids were allowed to rest for 15 s, back-side blotted for 8 s at 27°C and 95% humidity, and plunge-frozen in liquid ethane using an EM GP2 cryo-plunger (Leica). Frozen grids were stored in liquid nitrogen until FIB milling.

### 4.2 FIB milling for ion comparison

Lamellae were milled using argon, xenon, and gallium ion sources operated at 30 kV using a Hydra, Arctis, and Aquilos cryo-FIB-SEM instruments respectively (Thermo Fisher). Lamellae were prepared using *AutoTEM* (Thermo Fisher), see Table 1 for the currents used in each step of the protocol for each of the ion sources (Table 1). Stress relief cuts were performed, followed by rough milling to ~1 µm. Medium milling was performed using a cleaning cross-section pattern, followed by fine milling to a target thickness of ~500 nm. Final polishing was performed at ≤30 pA to a target thickness of 200 nm. SEM imaging was not performed during lamella preparation. Grids used for ion comparison were frozen in the same batch.

**Table 1:** Currents utilized for each ion during various milling steps.

| Milling step; Ion | Argon | Gallium | Xenon |
| --- | --- | --- | --- |
| Stress Relief | 0.74 nA | 0.3 nA | 0.3 nA |
| Rough Milling | 0.74 nA | 0.3 nA | 0.3 nA |
| Medium Milling | 0.2 nA | 0.1 nA | 0.1 nA |
| Fine Milling | 60 pA | 0.1 nA | 0.1 nA |
| Polish | 20 pA | 30 pA | 30 pA |

### 4.3 Nilas and 30 kV comparison

Lamellae were generated using an Aquilos2 cryo-FIB-SEM (Thermo Fisher Scientific) with a gallium liquid metal ion source, aligned at 30 kV, 8 kV, and 2 kV at the polishing current for each before milling. Grids were sputter coated (15 s, 30 mA and 0.10 mbar) with metallic platinum, GIS coated with organo-platinum for 2 min (1–2 µm as measured after rough milling by SEM); (Supplementary Fig. S4d), then sputter coated again with the same parameters. Lamellae sites were preferentially chosen in yeast clumps running perpendicular to the ion beam for a flat milling surface and with a width of ~5 µm. Automated milling to ~500 nm was performed in AutoTEM with a target milling angle of 20°to accommodate undertilt during the 2 kV polishing step. Stress relief cuts and rough milling were performed at 0.5 nA using a rectangular box. Medium milling steps were performed at 0.3 nA, 0.1 nA, and 50 pA with ±1.5°over/undertilt to protect the GIS layer, using a cleaning cross-section pattern. Lamellae were then manually thinned to ~300 nm at 50 pA with 0°or ±0.5°overtilt (Supplementary Fig. S4). The voltage was lowered to 8 kV and lamellae were thinned to ~200 nm using 27 pA with a cleaning cross-section pattern, with boxes drawn farther from the final lamella than typical to retain the GIS layer (Supplementary Fig. S4b). The milling z-depth was limited to 2 µm so that the intactness of the lamellae could be frequently monitored with FIB imaging. SEM imaging was used to verify GIS layer integrity after the 8 kV polishing step. This step typically resulted in the rejection of 15–20 percent of lamellae. The accelerating voltage was then lowered to 2 kV and the sample was overtilted by 10°. A rectangular pattern over the entire lamella face was used to mill (27 pA, 100 ns dwell time, and 85% beam x/y overlap, z-depth of ~0.005 µm) (Supplementary Fig. S4b,c). The sample was then undertilted by 10°and the same milling was performed. SEM imaging was used sparingly to check lamella integrity, with a maximum of five images at 15,000 x (or lower) magnification, 2 kV with 1 µs (or lower) dwell time to minimize electron damage (Supplementary Fig. S4d). Lamellae were not imaged with a higher voltage FIB following low voltage polishing. Aberrations are pronounced during the 2 kV under-tilt polishing step, particularly in higher tilts for lamellae closest to the ion beam. We attribute these aberrations to co-illumination of the metallic clip by the FIB. They can be mitigated with a steeper initial milling angle of 20 degrees, positioning lamellae higher on the grid, and increasing the magnification (to *>*20,000x) during polishing (Supplementary Fig. S4d,e). The 30 kV-milled comparison lamellae were generated on the same grid as Nilas lamellae and had the same GIS coating and milling angle. Briefly, stress relief cuts and rough milling were performed at 0.5 nA using a rectangle box. Multiple medium milling steps were performed at 0.3 nA, 0.1 nA, and 50 pA with ±1.5°over/undertilt to protect the GIS layer, using a cleaning cross-section pattern. Lamellae were then manually thinned to ~300 nm at 50 pA with no over/undertilt and to 100–200 nm with 30 pA.

### 4.4. Cryo-EM data collection and pre-processing

Cryo-EM data for ion comparison experiments were collected following the protocol described in ^19^ using a Thermo Fisher Krios 300 kV transmission electron microscope equipped with a Falcon 4i camera and Selectris X energy filter (Thermo Fisher) to a nominal magnification of 81,000× (pixel size of 0.93 Å) and a 100 µm objective aperture. Movies were collected to a total fluence of 50 e*^−^*/Å^2^ with 1800 frames in EER mode targeting a defocus of 0.5 µm. Cytoplasm was targeted by selecting regions of interest in a low-magnification overview and visually verified in final images. For the comparison of Nilas with 30 kV-milled lamellae, images were collected on a Thermo Fisher Krios G3 300 kV transmission electron microscope equipped with a K3 detector and energy filter (Gatan) at a nominal magnification of 81,000x, corresponding to a pixel size of 1.059 Å. Movies were collected to a total fluence of 50 e*^−^*/Å^2^ with 1 e*^−^*/Å^2^ per frame and a target defocus of 1 µm.

For all data, movie frames were aligned using MotionCor2^66^ and dose-weighted according to ^67^. CTF and thickness estimation were performed using CTFFIND5^68^. Additional cryo-EM data was collected using DeCo-LACE^52^ on a ThermoFisher Krios G3 300 kV transmission electron microscope. Movies were collected to a total fluence of 50 e*^−^*/Å^2^ with 1 e*^−^*/Å^2^ per frame at a target defocus of 1.2 µm with a beam size radius set to 221 nm. Movie pre-processing was performed as described in ^52^. Frames were aligned using MotionCor3^66^ and dose-weighted according to ^67^. CTF and thickness estimation were performed using CTFFIND5^68^.

### 4.5 2DTM of LSU

For the 30 kV ion comparison, a PDB model of the 80S ribosome (6Q8Y) ^25^ was used to generate a LSU template with Simulator ^69^ and a pixel size of 0.93 Å with 0 per atom *B*-factor scaling and 80 *B*-factor added to all atoms. 2DTM was performed using the programs match_template and refine_template *cis*TEM ^70^ as described in ^17^. All 2DTM in the main figures was performed using Leopard-EM ^31^. Briefly, we used a pixel size of 0.936 Å or 1.059 Å depending on the microscope parameters used for the dataset being processed. We simulated a 512 512 512 pixel template using the program ttsim3d with no re-centering, additional *B*-factor or hydrogen atoms and a PDB *B*-scaling of 0.5. We ran match_template in Leopard-EM ^31^ with uniform angular sampling, with a psi step of 1.5° and theta step 2.5° with C1 symmetry. The defocus search range was calculated to extend beyond the image defocus range in 200 Å steps. Peaks were extracted from the z-score map using a threshold of one false-positive per micrograph and were refined using refine_template at a defocus search of +/-100 Å in 20 Å steps and angular steps of 0.05°. Data were filtered to exclude peaks within 10 pixels and peaks that were not within the beam area. Optimize template ^31^ was used to find optimal pixel size. All molecular visualization was performed using ChimeraX by overlaying templates on images at the locations and orientations identified by 2DTM ^71^.

### 4.6 LSU Damage analysis

Ion comparison: Images were selected for damage analysis based on a *CTFFIND*5^68^ cross-correlation score *>*0.2 and thickness *>*140 nm and *<*220 nm, and were visually inspected to confirm cytoplasmic regions. Only images with *>*100 detected LSUs were included. CTFFIND5^68^ was used to estimate the lamella thickness where images were taken and was independently validated by the Beer-Lambert association between thickness and transmitted signal ln(*I/Io*) ^72^ (Supplementary Fig. S5e). For argon, gallium and xenon there were 58, 42 and 33 total images analyzed respectively. The protocol for quantifying damage was performed as described in ^9^. Detected particles were coordinate-transformed into lamella coordinates and binned in 10 nm depth bins from the lamella edge using particle coordinates from 2DTM and thickness, tilt-angle and tilt-axis from CTFFIND5. On a per-image basis, each particle z-score was normalized to the mean z-score of all particles in bins*>*70 nm from the lamella edge. Gaussians were fit to normalized z-score histograms for each 10 nm bin and the mean and standard deviation of the normalized z-score was estimated from these fits. The mean and standard deviation was plotted as a function of depth from the lamella surface and an exponential decay model was fit to the resulting curves excluding the 0–10 nm bins. Additionally, ribosomes were counted in 10 nm bins from the lamella surface and normalized to bins at a depth of *>*70 nm on a per image basis. Nilas and 30 kV comparison: Analysis was performed as described above with the following modifications. The least damaged Nilas lamellae were thinner than 160 nm, reflecting complete removal of the 8 kV damage layer by 2 kV polishing. Therefore, only Nilas images thinner than 160 nm were included in the damage analysis. 30 kV images were filtered to be *>*140 nm and *<*180 nm thick. These filters resulted in 97 Nilas-and 55 30 kV-milled lamella images. Because the Nilas filter excluded the 70 nm normalization bin, particle scores were normalized to bins *>*40 nm from the lamella surface. Exponential models were fit to the Gaussian parameters, excluding bins with n*<*150 LSUs which excluded the [0, 10] nm depth bin for the 30 kV condition. Relative signal plots were produced as done previously in ^9^ using electron scattering curves from ^28^. Ribosomes were counted in 10 nm bins from the lamella surface and normalized to bins at a depth of *>*70 nm from the image surface for gallium 30 kV and *>*40 nm from the image surface for Nilas on a per image basis. For (Supplemental Fig. S4c), 8 kV images were taken from uncurtained regions of the +/-4° lamella and filtered to be *>*160 nm and *<*200 nm thick to ensure they were representative of the 8 kV polished region (n=23 images). Images were normalized to bins at a depth of *>*70 nm from the surface.

### 4.7 3D Reconstructions

We truncated the LSU ribosome template (6Q8Y) by removing equivalent molecular weights of RNA and protein to 1 MDa. We selected the 40 images from each condition with the highest mean of the top-10 full LSU 2DTM z-scores and searched them using our truncated template. We found more particles in 2 kV images (3,113) so randomly sub-selected to 2,748 particles from each condition. Micrographs and particles were imported into cryoSPARC^73^ using custom Python scripts and the 3D reconstruction was performed using homogeneous refinement with the particle positions, orientations and defoci from 2DTM without further refinement. ResLog in cryoSPARC was used to generate Rosenthal-Henderson B-factor plots ^30,74^. Homogeneous refinement half maps were used to estimate local resolution in Relion using local-resolution estimation ^75,76^ with a box size of 12 pixels. Resulting volumes, and the isolated non-template region of the LSU, were visualized using ChimeraX at the same level (0.28) and colored by local resolution ^71^. The per voxel local resolution of the non-template region of the LSU was filtered to resolutions lower than 15 Å and plotted for each condition. A spatial block bootstrap significance test was utilized to account for the non-independence of neighboring local-resolution measurements. Non-overlapping voxel blocks (12 pixels) were sampled with replacement 10,000 times to estimate the mean difference between conditions. The difference was considered significant if the 95% confidence interval excludes 0. Using homogeneous refinement, the isolated non-template region was masked to determine its global FSC for both conditions.

### 4.8 2DTM of SSU and Truncated Ribosome Search

The SSU body was generated from (PDB:6Q8Y) as described in ^31^ and 2DTM was performed as described above. The LSU ribosome template (PDB:6Q8Y) was truncated by removing equivalent molecular weights of RNA and protein to produce a template of 300 kDa, 500 kDa, 750 kDa and 1000 kDa. The ten images containing the highest mean z-score of the top 10 LSUs were selected for both Nilas-milled and 30 kV-milled lamellae (Fig. 2g) and template matching performed with Leopard-EM with each truncated template. The optimal pixel size was utilized for for these searches and found by using optimize_template. The 2DTM z-scores of the top 10 particles and concentration of particles above the detection threshold were plotted for templates with data at all 10 images. Concentration was calculated using the volume calculated using the thickness of lamellae at each image from CTFFIND5 and the beam size. The 2DTM z-score threshold was calculated for different numbers of cross-correlations using the equation:

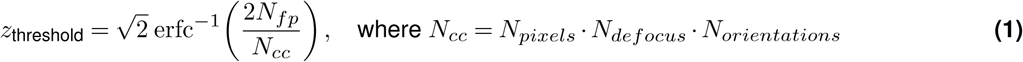

Where erfc is the complimentary error function, *N_fp_* is the expected number of false positives (set to 1), *N_pixels_*is the number of pixels searched, *N_defocus_* is the number of defocus planes searched, and *N_orientations_* is the number of orientations searched. (*N_defocus_*) is adjusted for lamellae of different thicknesses and (*N_orientations_*) can be reduced for templates with symmetry.

### 4.9 2DTM of Nuclear targets

Ten images of thin nuclear regions from Nilas and 30 kV-milled lamellae were identified and the RNA polymerase III template and proteasome (PDB: 8BWS^39^, PDB: 1RYP, ^35^) were simulated and used as templates for a 2DTM search in Leopard-EM. Briefly, we simulated a 512 512 512 pixel template using the program ttsim3d with no re-centering, additional *B*-factor or hydrogen atoms and a PDB *B*-scaling of 0.5 and ran match_template in Leopard-EM ^31^ with uniform angular sampling, with a psi step of 1.5°and theta step 2.5°. Template pixel size was optimized using optimize_template and match_template was re-run at the optimal pixel size. We used the same refine_template parameters as for the LSU 2DTM and we excluded peaks beneath a z-score threshold of 1 false positive per 10 images. Additionally, images were filtered to only include findings within the illuminated beam area and to remove peaks closer than 10 pixels. All images were visually inspected and peaks over ice contamination were eliminated. For the insets in (Supplemental Fig. S9a,c) images were Wiener filtered (3×3 window) and low pass filtered (sigma =1) for visualization. For DNA visualization insets in in (Supplemental Fig. S9b), images were Wiener filtered (5×5 window) and Gaussian filtered (sigma =1) for visualization.

### 4.10 2DTM of Cytoplasmic targets

The ten cytoplasmic images with the highest mean 2DTM z-score of the top 10 LSUs from Nilas-and 30 kV-milled lamellae were searched for the following cytoplasmic templates: fatty acid synthase (PDB: 2UV8, biological assembly 1) ^32^, phosphofructokinase (PDB: 3O8O) ^37^ and pyruvate kinase (PDB: 1A3X) ^41^ using Leopard-EM with the same parameters as for the nuclear templates and applying symmetry as indicated in (Fig. 5). The 2DTM z-score threshold was determined by 1 false positive per 10 images. Additionally, images were filtered to only include findings within the illuminated beam area and to remove peaks closer than 10 pixels. All images were visually inspected and peaks over ice contamination were eliminated. For 1A3X, the PDB file contained 2/4 chains and was duplicated using symmetry to reconstruct the full tetramer. Results were visualized using ChimeraX and images were Wiener filtered (3×3 window) and low pass filtered (sigma =1) for visualization.

### 4.11 SRIM simulations

Frozen hydrated yeast cell matter was approximated as 0.14 parts carbon 0.74 percent oxygen and 0.10 percent hydrogen and given a density of 1.1 g/cm3^34,77^. All SRIM simulations were run with 5000 ions. Angles in SRIM are defined as relative to the surface normal. Therefore, milling was simulated with

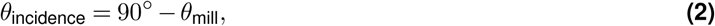

with the exception of 0*^◦^* milling, which was simulated at *θ*_incidence_ = 88*^◦^* rather than 90*^◦^* so that ions would penetrate into sample. Ion ranges were exported from SRIM output Range.txt and plotted with Python ^26^.

### 4.12 Measuring 2 kV region

The extent of 2 kV polishing was determined visually by the characteristic curtaining pattern and increased intensity of thinner regions produced in cellular material. In 10°samples, the 2 kV polish extended to the cell boundary, which will limit the measurement. Images from the 2 kV and 8 kV regions were analyzed using the LSU damage analysis described above.

### 4.13 Calculation of expected number of macromolecules in cytoplasm and nucleus

The expected number of macromolecules per imaging volume was estimated using whole organism proteomics data compiled from multiple datasets on paxdb ^33^. The following equation was used in order to determine the expected subunits per cell.

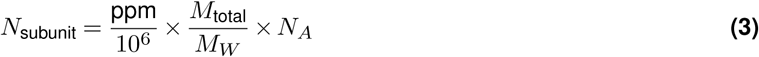

where ppm is the paxdb abundance value (mg protein per kg total cellular protein), *M*_total_ is the total protein mass per cell (6 10*^−^*^12^ g), *M_W_* is the monomer molecular weight (g mol*^−^*^1^), and *N_A_* is Avogadro’s number (6.022 10^23^ mol*^−^*^1^).

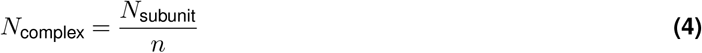

where *n* is the number of copies of the subunit per complex.

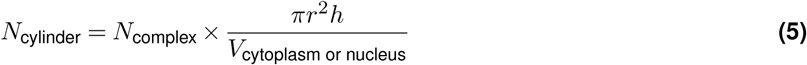

where *r* is the beam radius, *h* is the lamella average thickness for the Nilas-and 30 kV-milled conditions respectively, and *V* is the volume of the cytoplasm (*V*_cytoplasm_ = 27 *µ*m^3^) and nucleus (*V*_nucleus_ = 2 *µ*m^3^), excluding the nucleolus ^34^.

### 4.14 Probability of false-positive calculation

The false-positive probability of individual particle detections was estimated using the Gaussian background noise model

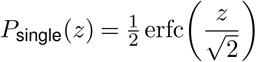

where *P*_single_(*z*) is the probability that the cross-correlation z-score at a single search location is a false-positive, erfc is the complementary error function, and *z* is z-score of the peak, as was done by ^7,31^.

### 4.13 Idealized damage curve without subsurface damage

To estimate damage that is only affected by LSU ablation and not subsurface damage, LSUs were approximated as uniform density spheres with a radius of 15 nm with even distribution across 10 nm bins. Any region of the sphere lying outside the lamella was subtracted from the LSUs volume, and the z-score was adjusted according to

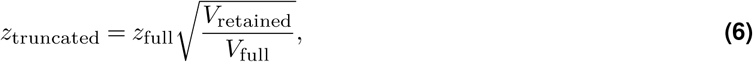

where *z_truncated_*is the 2DTM z-score of the truncated LSU, *z_full_* is the 2DTM z-score of a full LSU, *V_retained_* is the volume of the sphere lying inside the lamella, and *V*_full_ is the volume of the full sphere.

## A. Code and data availability

The code utilized for the FIB-milling damage analysis can be found on the Lucas lab gitub page at https://github.com/Lucaslab-Berkeley/2DTM_FIB_Damage_analysis. Data collected for this manuscript will be published in EMPIAR before publication.

## Acknowledgments

The authors would like to thank Joshua Dickerson for invaluable project guidance, critical feedback on the manuscript and assistance setting up DeCo-LACE data collection and implementing image processing pipelines ^52^. We thank Lucille A. Giannuzzi, Joseph Michael and Reed Yalisove for thoughtful comments and advice in applying developed material science protocols to biological specimens. Matthew Giammar, Wolfram Seifert Davila and Kithimini Herath graciously shared scripts for result visualization and Leopard-EM to cryoSPARC particle stack conversion. The authors would like to thank scientists in the cryo-EM facilities utilized in this research including Elizabeth Montana and Garrett Greenan from Biohub, David Bulkeley and Glenn Gilbert from UCSF, and Lydia Marie-Joubert from SLAC for practical support and guidance. The plasma FIB experiments were performed as part of the Biohub Imaging Residency Program, and much of the data processing was preformed using Biohub computational resources. We thank Bridget Carragher, David Agard and Clint Potter for the use of resources and their guidance and practical support. This project has been made possible in part by grant number 2025-358053 from the Chan Zuckerberg Initiative DAF, an advised fund of Silicon Valley Community Foundation and by support from an NIH Director’s New Innovator Award (DP2GM159184). BAL is a Searle Scholar and a Shurl and Kay Curci Scholar and LNH is supported by the NSF GRFP.

## Author contributions

L.N.H conceptualization, validation, methodology, formal analysis, investigation, data curation, writing-original draft, writing-review and editing, visualization. B.A.L conceptualization, validation, data curation, writing - original draft, writing-review and editing, supervision, project administration, funding acquisition. J.P and P.N. software.

## Supplementary Information

**Figure S1.**
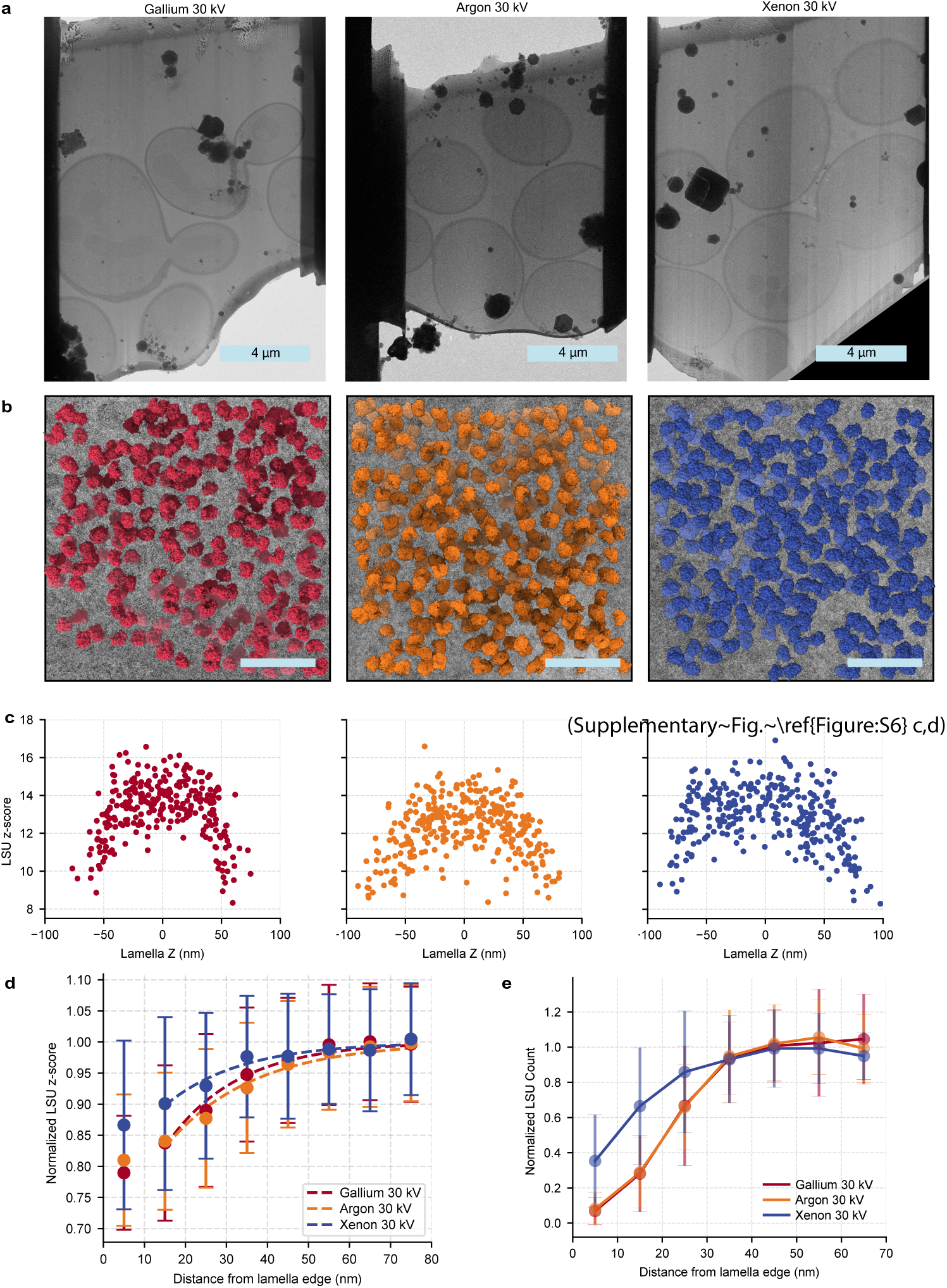
Xenon reduces FIB Damage at 30 kV. **(a)** TEM images of example lamellae milled with gallium, argon and xenon at 30 kV. Scale bar: 4 µm **(b)** Example images from lamellae milled with each ion overlaid with ribosomal LSUs in the locations and orientations identified with 2DTM. Scale bar: 50 nm. **(c)** Scatterplot of the mean 2DTM z-scores of LSUs as a function of distance from lamella edge in 10 nm bins normalized to undamaged bins in the center of the lamella from the same images of argon-milled (yellow), xenon-milled (blue) or gallium-milled (red) lamellae. **(d)** Scatterplot showing the normalized 2DTM z-scores of multiple images, plotting SNR as a function of distance from lamella edge. All z-scores were normalized to the undamaged bins > 70 nm from the lamella edge from the same image. Curves represent the fits of the exponential decay function. Error bars indicate the standard deviation. **(e)** Number of LSUs identified in depth bins from lamella surface from the same subset of images from part (d). Normalized to full undamaged bin counts at > 70 nm from the lamella edge in individual images. Error bars indicate the standard deviation.

**Figure S2.**
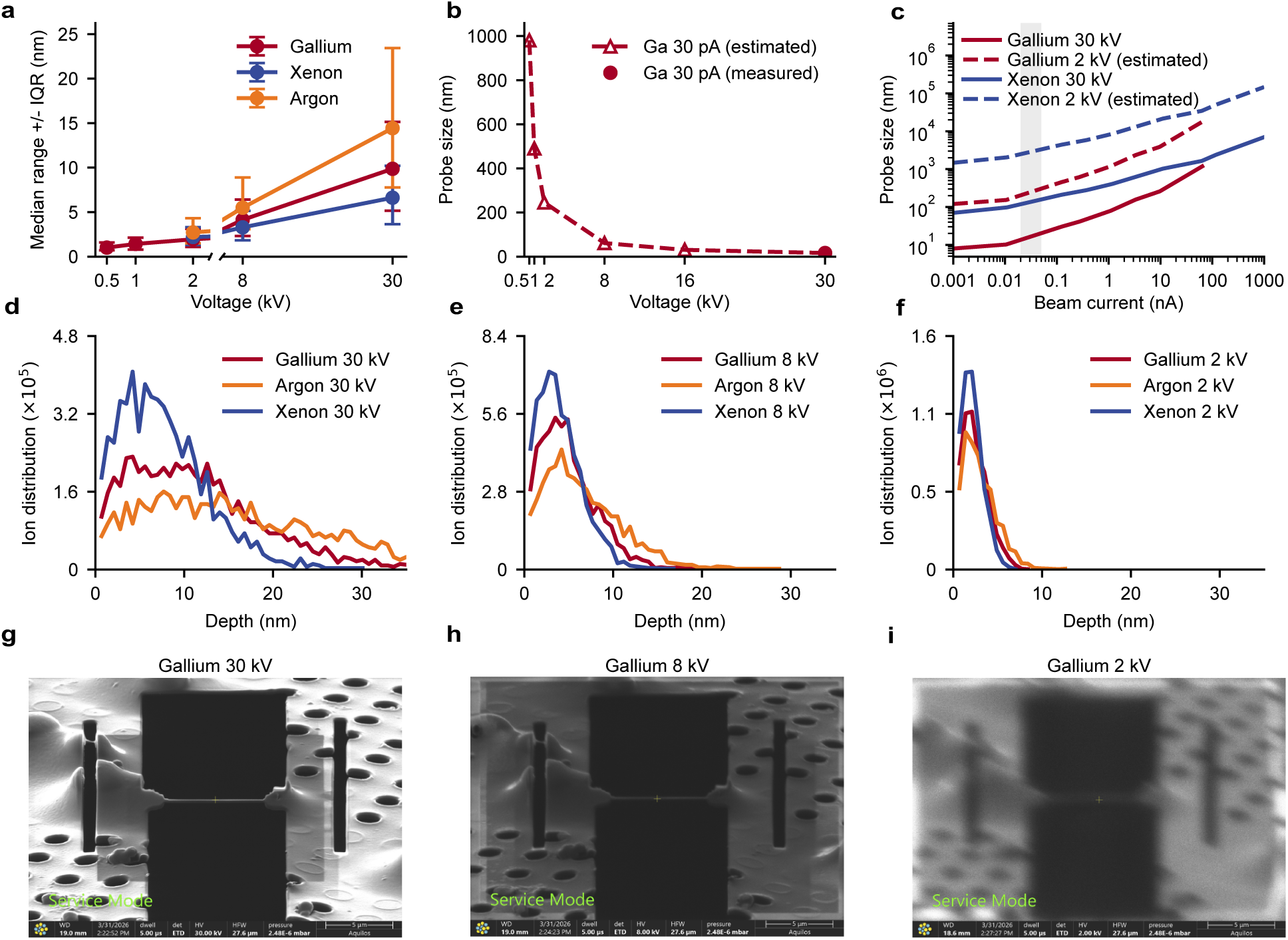
Justifying the selection of gallium for low-voltage milling. **(a)** SRIM simulated median range for gallium, argon and xenon at voltages from 0.5 kV to 30 kV. Error bars indicate the IQR. **(b)** Probe size of Gallium at voltages from 0.5 kV to 30 kV. Value for the 30 kV probe was experimentally determined ^1^ and lower voltage probe sizes were predicted using 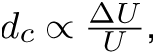 where *dc* is the probe diameter, Δ*U* is the energy spread of the beam, and *U* is the accelerating voltage^2^. **(c)** Probe size for Gallium and Xenon across beam currents, produced as in (b). Grey highlighted region indicates common polishing currents. **(d,e,g)** SRIM simulation for ion distribution of Argon, Gallium Xenon at 30 kV, 8 kV and 2 kV in “frozen yeast” (see methods). **(g,h,i)** FIB images at 30 kV, 8 kV, and 2 kV showing image blurring due to increased probe size at low voltage.

**Figure S3.**
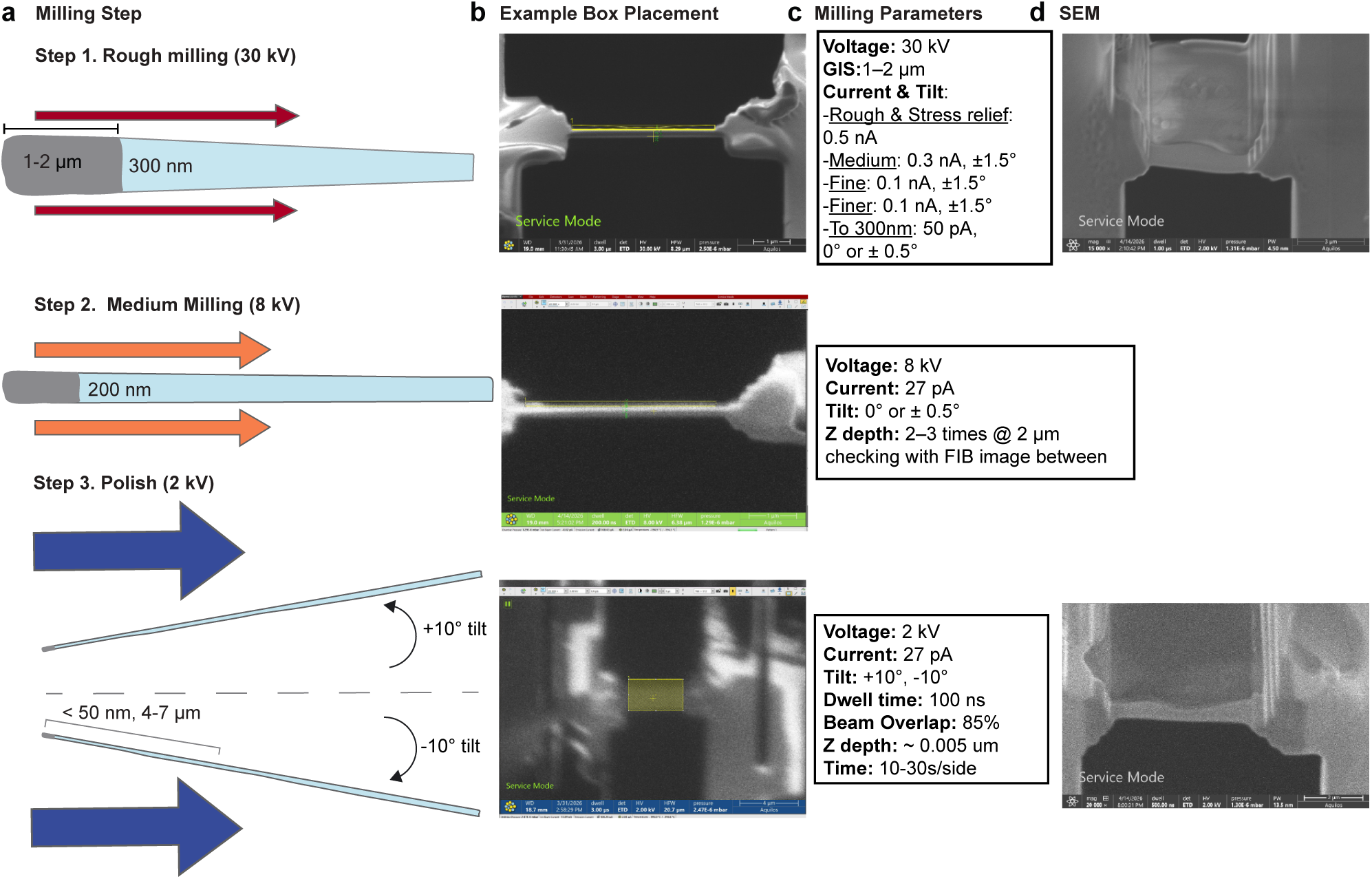
Steps, example pattern placements, and milling parameters for Nilas. **(a)** Schematic of the three voltages steps of the Nilas method. **(b)** Example milling pattern placement for each Nilas step as seen in the FIB image. Cleaning cross section patterns are placed as typical for 30 kV and without overlaying the visible area of the lamellae due to the increased beam size at 8 kV. For 2 kV a rectangle pattern is drawn broadly across the lamella surface. **(c)** Milling parameters used at each Nilas step. **(d)** SEM images of lamellae after completion of the corresponding step, SEM contrast changes as lamella is thinned.

**Figure S4.**
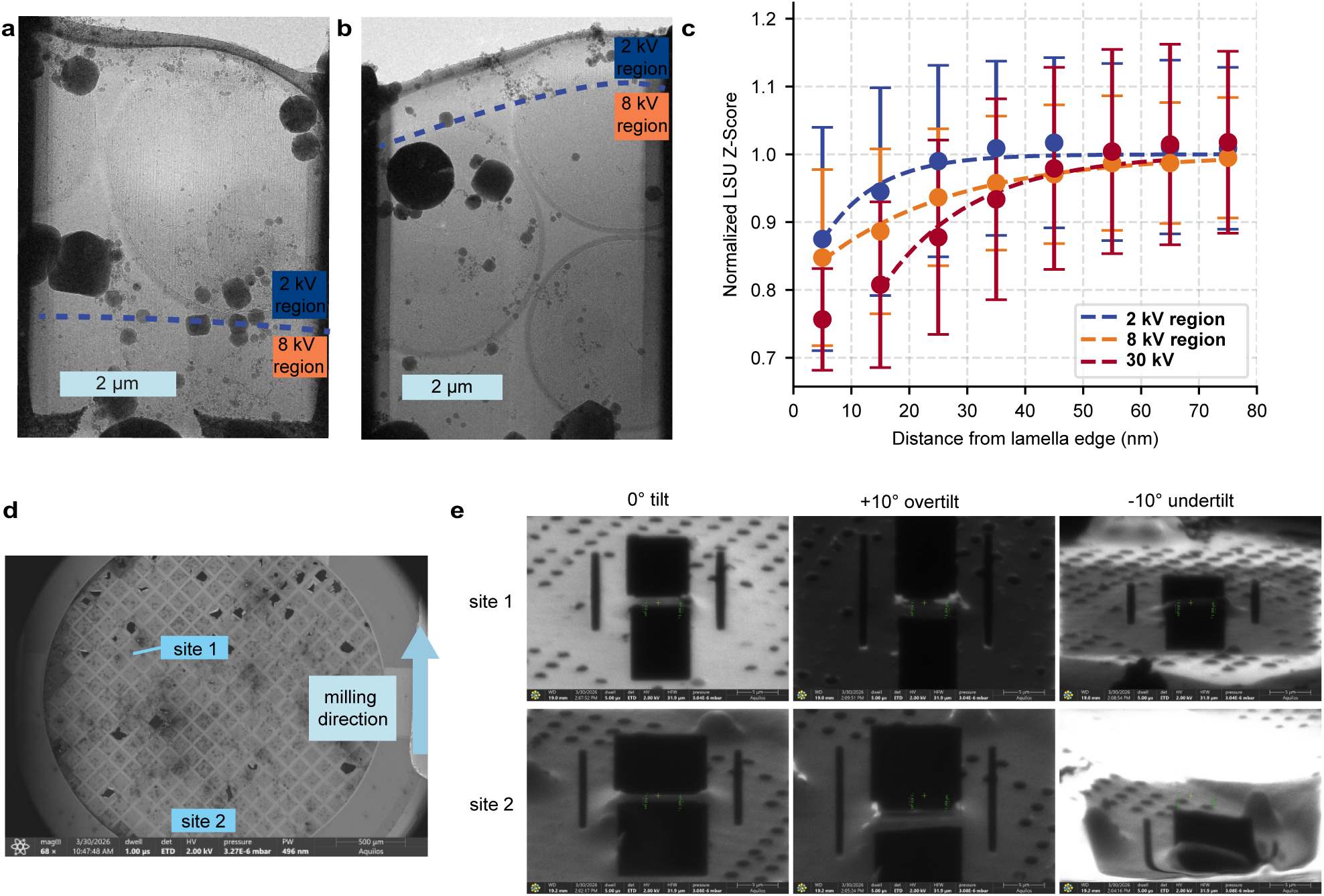
A ±10 degree tilt is required to extend 2 kV damage region. **(a)** Example TEM image of a lamella polished at 2 kV with +10 degree tilt (relative to initial milling angle). Scale bar: 2 µm. **(b)** Example TEM image of a lamella polished at 2 kV with a +4 degree tilt. Scale bar: 2 µm. **(c)** Scatterplot of the mean 2DTM z-scores of LSUs binned by distance to lamella edge normalized to undamaged bins in the center of images in gallium-milled at 30 kV (red), gallium-milled at 8 kV (orange) or gallium-milled at 2 kV (red) lamellae regions. Curves represent fits of the exponential decay function. Error bars indicate the standard deviation. (**d**) An SEM image showing the overview of a grid with two milling sites: site 1 (nearer to the FIB source) and site 2 (further from the FIB source). The FIB source is towards the bottom of the image. **(e)** Images depicting the 2 kV FIB image warping during the 10 degree under-tilt as a result of the ion beam illuminating the clip ring.

**Figure S5.**
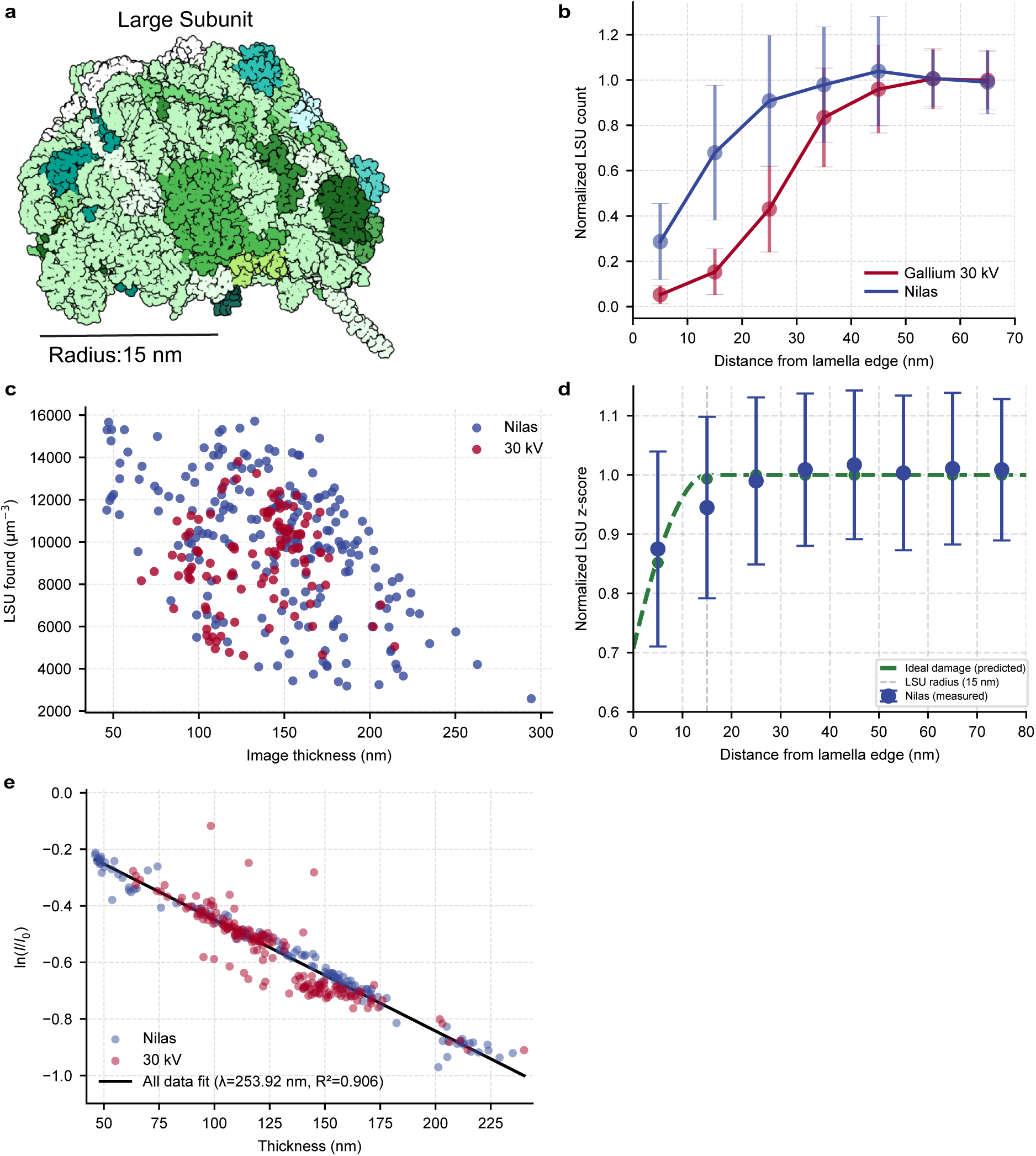
Measuring FIB-damage with 2DTM of the LSU. **(a)** A model of the LSU of the ribosome (PDB:6Q8Y) indicating its radius of 15 nm. **(b)** The number of LSUs identified in 10 nm bins relative to the lamella surface normalized to the number of significant LSU detections in an undamaged bin. Error bars indicate the standard deviation. **(c)** The concentration of LSUs detected per image in Nilas and 30 kV images as a function of thicknesses. **(d)** The predicted ideal damage curve (green) is a result of the effect of LSU ablation on the 2DTM z-score ignoring the effects of sub-surface damage. Nilas-milled lamellae damage curve (blue) from (Fig. 2e) for comparison. **(e)** The CTFFIND5 determined thickness plotted against the natural log of image intensity over vacuum intensity. The plot demonstrates a Beer-Lambert law relationship between thickness and transmitted signal ln(*I/I*_0_).

**Figure S6.**
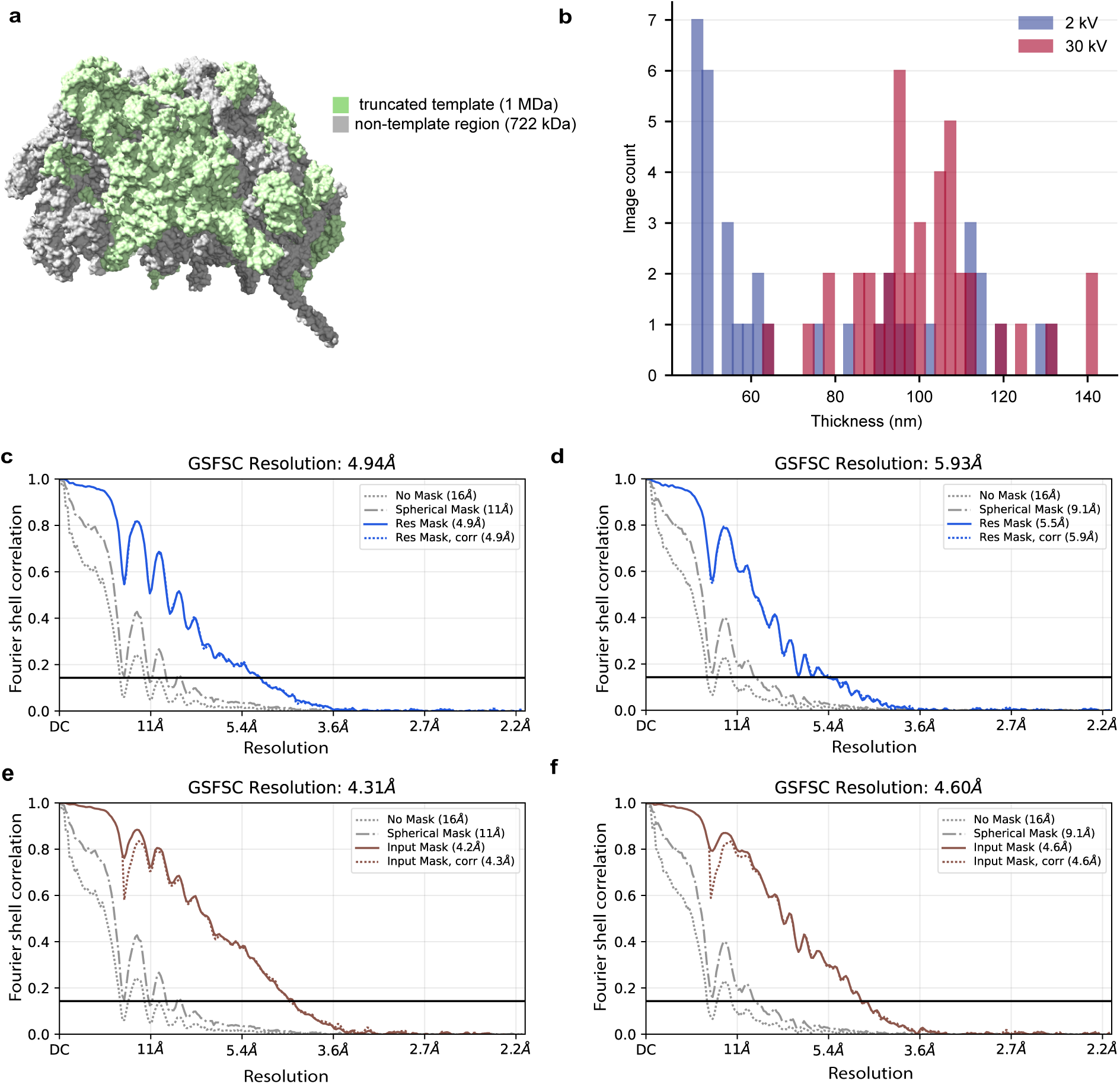
Baited reconstructions using 1 MDa LSU fragment. **(a)** LSU truncated 1 MDa template in grey and non-template region (722 kDA) in green. **(b)** Histogram of CTFFIND5 estimated thickness for images used in the reconstructions in (Fig. 3a) for each condition. **(c)** Fourier shell correlation (FSC) curve for the Nilas homogeneous reconstruction in (Fig. 3a). FSC oscillation is due to narrow defocus range of 2DTM-identified particles. Resolution determined at FSC value of 0.143 (black line). **(d)** FSC curve for the 30 kV LSU homogeneous reconstruction in (Fig. 3a) FSC oscillation is due to narrow defocus range of 2DTM-identified particles. Resolution determined at FSC value of 0.143 (black line). **(e)** FSC curve for the LSU region omitted from the 2DTM template from the homogeneous reconstruction using particles identified in 30 kV-milled lamellae in (Fig. 3c). **(f)** As for (e) using particles identified in Nilas-milled lamellae.

**Figure S7.**
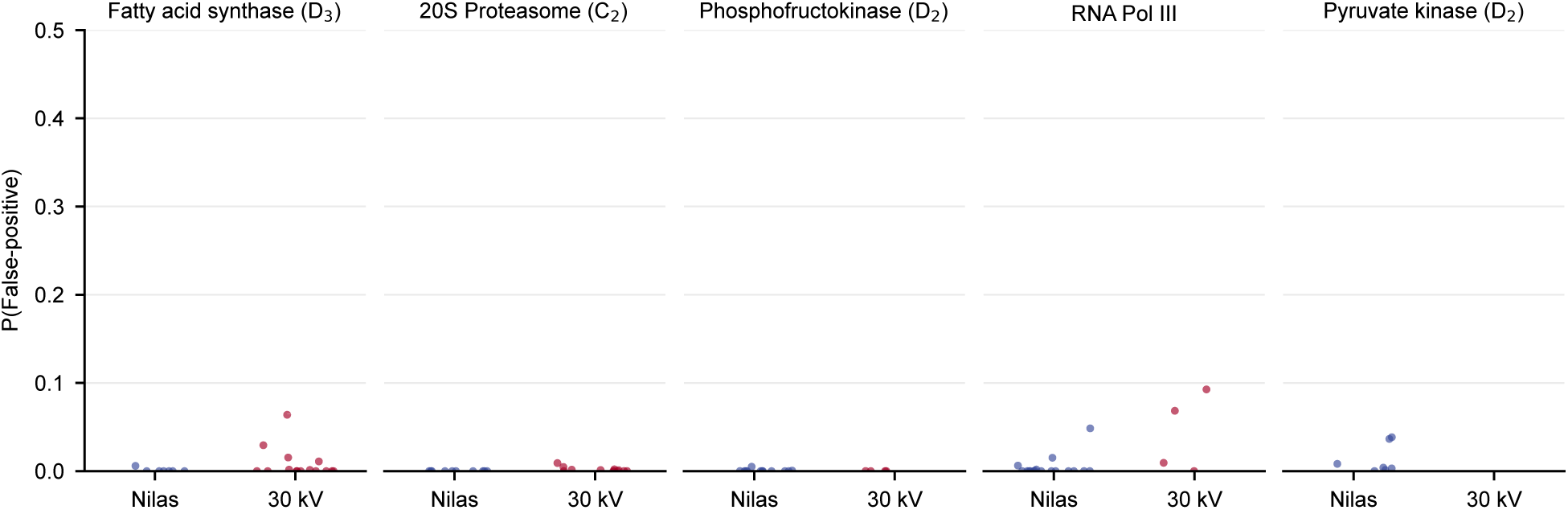
The probability of false positive detections with 2DTM on non-ribosomes. The false positive probability of 2DTM peaks based on their z-scores as calculated by the Gaussian noise background model for 2DTM peaks from (Fig 5).

**Figure S8.**
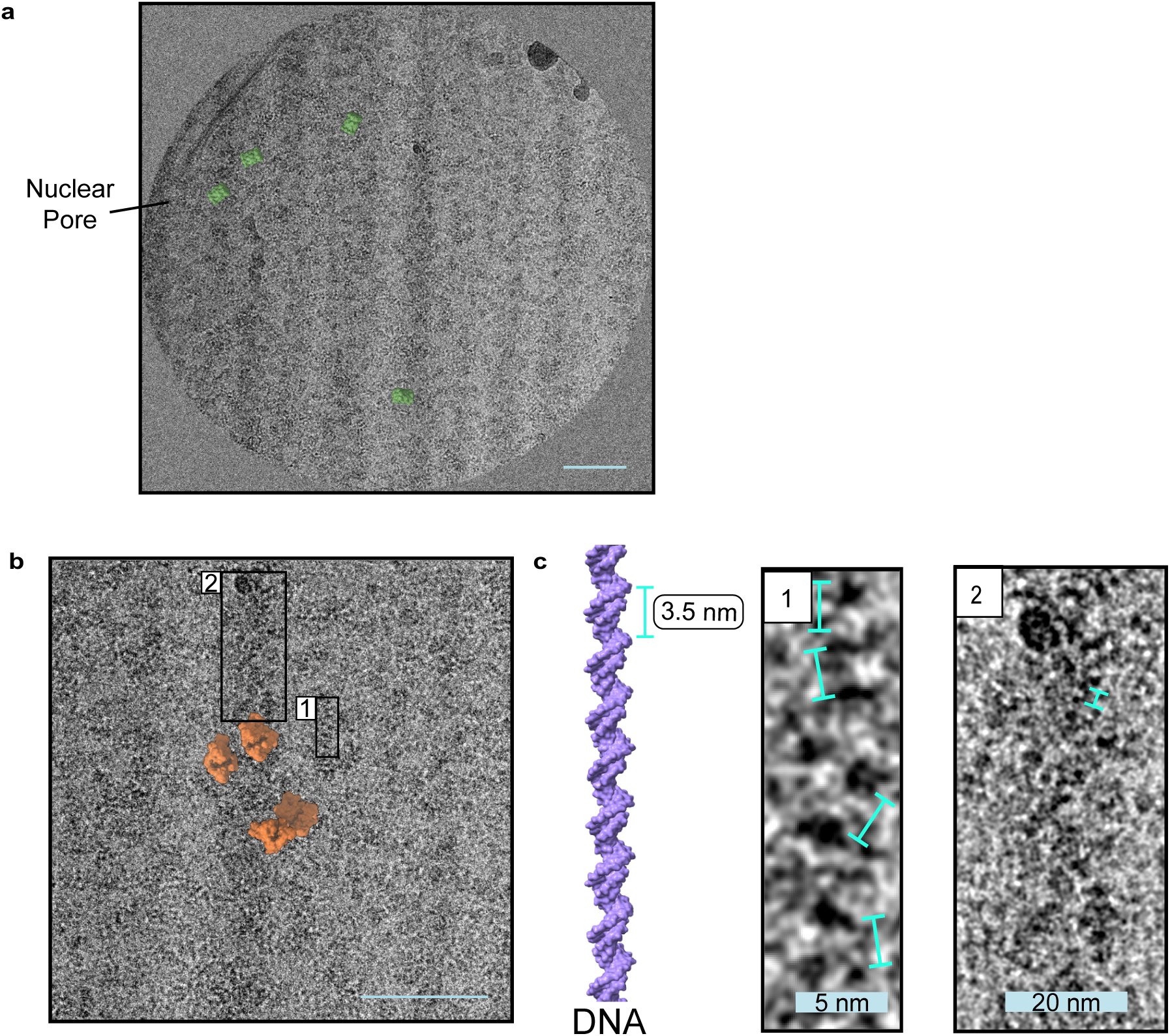
Observations enabled by thin lamellae with Nilas. (**a**) Overlay of the proteasome template for locations and orientations identified by 2DTM on an image region with nuclear membrane and nuclear pore visible. Scale bar: 50 nm. (**b**) Overlay of RNA polymerase III template for locations and orientations identified by 2DTM on an image region with RNA polymerase III in orange. Scale bar: 50 nm. (**c**) A DNA model (PDB: 4BNA) scaled to the same magnification as (1). Inset 1: from part (b) showing alternating contrast caused by DNA. Teal markers represent 3.5 nm which is consistent with the pitch of DNA. Scale bar: 5 nm. Inset 2: from part (b), suspected DNA super-coiling. Scale bar: 20 nm.

**Figure S9.**
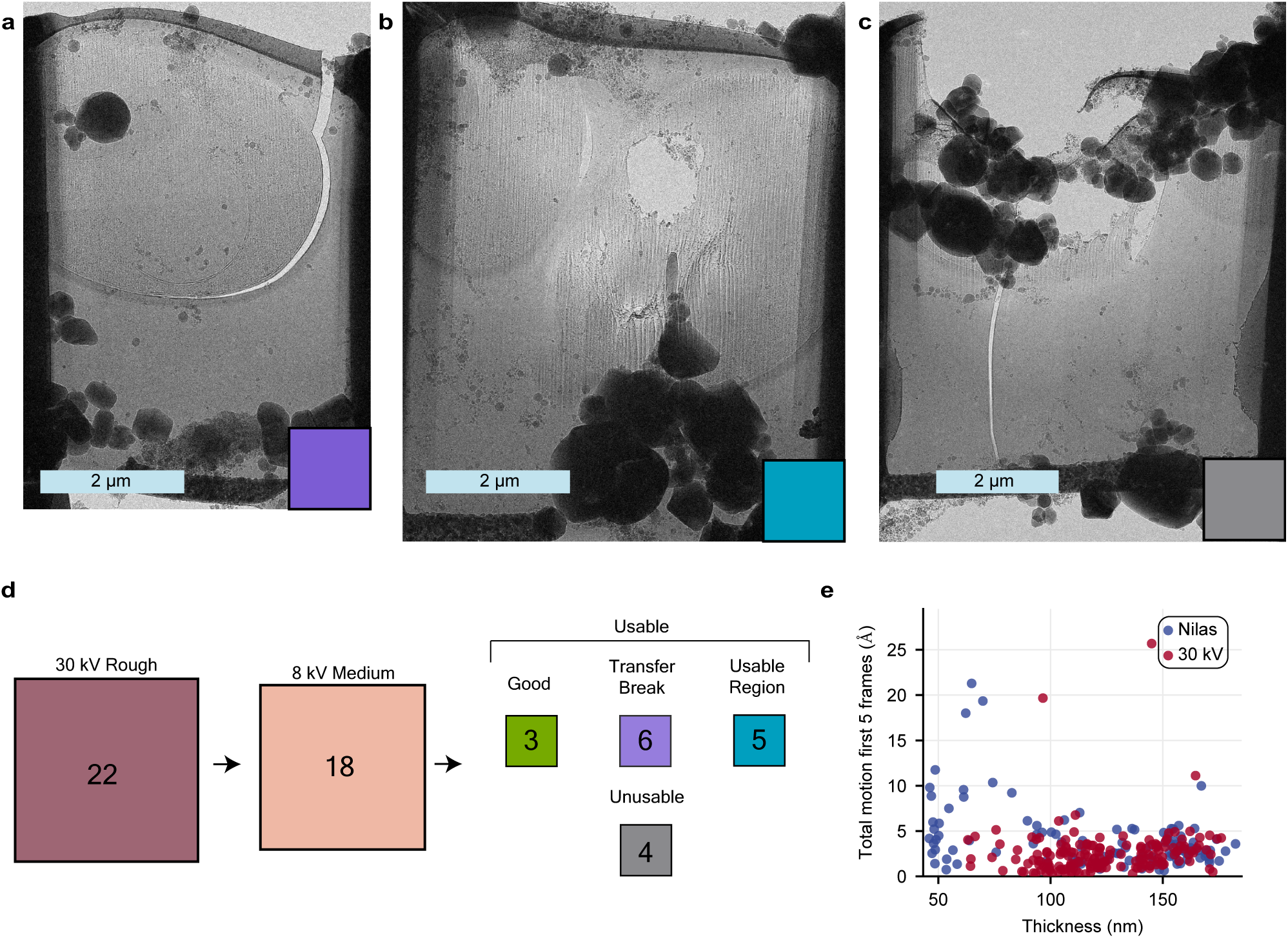
The throughput and stability of Nilas lamellae. TEM images of an example lamella that **(a)** broke at the cell membrane and cell wall boundary between the FIB and TEM position **(b)** was over-milled with 2 kV resulting in excessive curtaining and a hole in a less dense region and **(c)** had the organo-platinum layer milled away resulting in breakage. Scale bar: 2 µm. **(d)** The throughput of Nilas. 22 Nilas lamellae were originally attempted and 18 were successfully milled with 8 kV. Of those 18, 3 were excellent, 6 broke during transfer between the FIB and TEM (a), 5 had some usable regions but contained holes or cracks (b) and 4 were completely unusable (c). **(e)** Summed inter-frame global motion of the first 5 frames (5 e*^−^*/Å^2^) for Nilas and 30 kV-milled images at various thicknesses.

